# Zn-dependent structural transition of SOD1 modulates its ability to undergo liquid-liquid phase separation

**DOI:** 10.1101/2022.02.20.481199

**Authors:** Bidisha Das, Sumangal Roychowdhury, Priyesh Mohanty, Azamat Rizuan, Jeetain Mittal, Krishnananda Chattopadhyay

**Affiliations:** Structural Biology and Bioinformatics Division, CSIR- Indian Institute of Chemical Biology, Kolkata- 700 032, India; Academy of Scientific and Innovative Research (AcSIR), Ghaziabad, India; Artie McFerrin Department of Chemical Engineering, Texas A&M University, College Station, TX 78743

**Keywords:** Amyotrophic Lateral Sclerosis (ALS), SOD1, Liquid-liquid phase separation (LLPS), Aggregation, Molecular dynamics (MD) simulation

## Abstract

The toxic gain of function of Cu/Zn superoxide dismutase (SOD1) associated with the neurodegenerative disease - Amyotrophic lateral sclerosis (ALS), is believed to occur via misfolding and/or aggregation. SOD1 is also associated with stress granules (SGs) which are a type of membraneless organelle believed to form via liquid-liquid phase separation (LLPS) of several proteins containing low-complexity, disordered regions. Using a combination of experiments and computer simulations, we report here that structural disorder in two loop regions of SOD1 induced by the absence of metal cofactor - Zn, triggers its LLPS. The phase-separated droplets give rise to aggregates which eventually form toxic amyloids upon prolonged incubation. The addition of exogenous Zn to immature, metal-free SOD1 and the severe ALS mutant - I113T, stabilized the loops and restored the folded structure, thereby inhibiting LLPS and subsequent aggregation. In contrast, the Zn-induced inhibition of LLPS and aggregation was found to be partial in the case of another severe ALS-associated mutant - G85R, which exhibits reduced Zn-binding. Moreover, a less-severe ALS mutant - G37R with perturbed Cu binding does not undergo LLPS. In conclusion, our work establishes a role for Zn-dependent modulation of SOD1 disorder and LLPS as a precursor phenomenon which may lead to the formation of toxic amyloids associated with ALS.

**Significance Statement:** The formation of membraneless organelles such as stress granules (SGs) is believed to occur through the process of liquid-liquid phase separation (LLPS) and involves numerous proteins containing intrinsically disordered regions. Whether SOD1, which is also associated with SGs and whose aggregation is associated with Amyotrophic lateral sclerosis (ALS), can independently undergo LLPS, is not known. SOD1 is a metalloenzyme which is stabilized by the metal co-factor - Zn. In this work, we utilize experimental and simulation techniques to highlight the modulation of SOD1 LLPS propensity in a Zn-dependent manner due to underlying conformational transitions between folded and partially disordered states. Our work establishes a link between SOD1 LLPS and aggregation, which is relevant to ALS pathogenesis.

## Introduction

Amyotrophic lateral sclerosis (ALS) is an adult-onset progressive neurodegenerative disease that affects motor neurons of the brain and spinal cord, resulting in loss of control over voluntary muscle, paralysis, and death, often by respiratory failure (1). About 8-10% of ALS cases are familial (fALS) (2), while the rest occur sporadically (sALS). There are more than 140 point mutations of human Cu/Zn superoxide dismutase (SOD1), which are associated with fALS (3). Cu/Zn SOD1 is a highly conserved 153 amino acids long metalloenzyme, encoded by the SOD1 gene located on chromosome 21. It is the primary scavenger of reactive oxygen species, catalysing the conversion of superoxide radicals to hydrogen peroxide and water in a two-step reaction.

Each SOD1 monomer consists of eight anti-parallel β strands arranged in a β barrel structure. Extensive structural and biophysical experiments (4) have previously established that the immature, monomeric state of SOD1 contains two disordered regions (Fig. 1a-b, S1a-c), one from amino acids 49 to 83 and the other from 121 to 142. The first region (which corresponds to loop-IV) is referred to as the “Zn-binding” loop. The second region corresponds to loop-VII and is commonly referred to as the “electrostatic loop”. SOD1 matures through three post-translational modifications, namely the zinc (Zn) binding, copper (Cu) insertion via Cu chaperone protein (CCS), and disulfide bond formation (5–7). While Cu is responsible for the activity of the enzyme, Zn binding to loop IV presumably decreases the disorder by guarding the electrostatic loop and structuring the Cu insertion site (8). In the early steps of SOD1 biogenesis and its intracellular transport to mitochondrial inter-membrane space, the protein in its monomeric metal-free state is highly disordered in solution, and prone to misfolding and subsequent aggregation (9).

**Figure 1.**
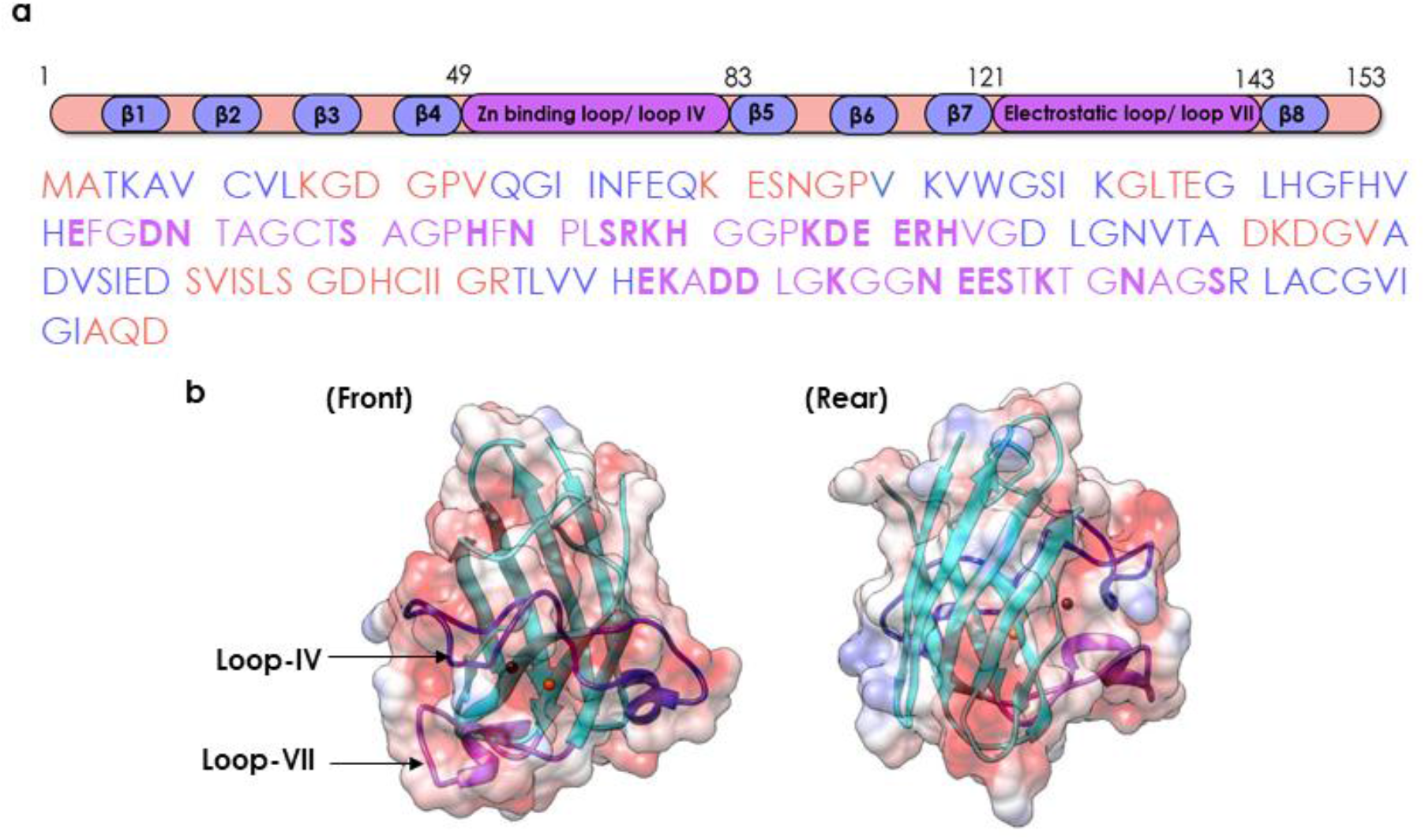
The structure of SOD1. **a)** Human SOD1 amino acid sequence comprised of 8 beta sheets (highlighted in blue) connected by 7 loop regions (in black); Zn binding loop IV highlighted in red; polar residues highlighted in bold within two loops. **b)** Cartoon representation of SOD1 monomer highlighting loops IV and VII (Coulombic surface coloring: red=negative, white=neutral, blue=positive).

The assembly and aggregation of SOD1 ALS mutants have been shown in cytosolic SGs (10–12). SGs are membraneless organelles composed of proteins and nucleic acids, possessing liquid-like properties (13). They originate as a response to intracellular stress through a process called LLPS and may revert to their non-droplet state once the stress is dissipated (14–18). The presence of SGs comprising predominantly of SOD1, TAR DNA Binding Protein 43 (TDP-43), and FUS form the central pathogenic hallmark of ALS (19, 20). Previous studies have demonstrated the phase separation of other intrinsically disordered proteins (IDPs) involved in the formation of SGs like FUS and TDP-43. It is hence important to investigate if SOD1, which is also a component of SGs, would phase separate under suitable conditions. It may also be noted that while a recent study reports that Zn promotes LLPS of tau protein (21), the phase behavior of metal-containing proteins has been relatively unexplored.

In this study, we observed that while the disulfide reduced, fully metalated form of SOD1 (WTSOD1^2SH^) does not undergo LLPS, the metal-free Apo variant (ApoSOD1^2SH^) forms liquid droplets, which can be reversed by Zn addition. We then show that two severe ALS mutants, namely I113T SOD12SH and G85R SOD1^2SH^ with destabilized Zn binding sites (and low affinity towards Zn) also undergo LLPS, while another mutant G37R SOD1^2SH^ (with less disease severity and strong binding to Zn) does not. For ApoSOD1^2SH^, I113T1SOD^2SH^ and G85RSOD1^2SH^ variants, LLPS is followed by aggregation – although, unlike the liquid droplets, the aggregated state could not be reversed by the addition of Zn. Interestingly, Cu, which is primarily responsible for dismutase activity, does not have any effect on LLPS and/or aggregation. Using Fluorescence Correlation Spectroscopy (FCS) and Fourier-transform infrared spectroscopy (FTIR), we find that the LLPS and subsequent aggregation is regulated by the conformational transition between a disordered and a relatively compact folded state. The extent of intrinsic disorder in various SOD1 monomeric states was further characterized using all-atom molecular dynamics (MD) simulations. A coarse-grained simulation of the ApoSOD1^2SH^ condensed phase suggested a stabilization of the condensed phase via both electrostatic and hydrophobic interactions. Overall, the cofactor Zn, by virtue of its efficient shielding (22) of loops IV and VII, acts as a switch to control conformational disorder and the phase separation propensity of SOD1.

## Results

SOD1 is different from other SG proteins that are implicated in ALS such as FUS and TDP-43 as it lacks long intrinsically disordered regions (IDRs) which are known to drive the LLPS of these other proteins. It is also much smaller (153 aa) than these other proteins (FUS: 526 aa, TDP-43: 414 aa) thereby reducing its ability to form comparable, multivalent interactions. As SOD1 is an integral component of SGs, it is important to determine whether it can undergo LLPS without the presence of other SG proteins. To answer this question, we conducted several biophysical measurements to check if SOD1 phase separates in vitro. All experiments mentioned in this paper were carried out using reduced monomeric protein variants (checked typically using mass spectrometry and gel electrophoresis as shown in fig. S1d-e) using 20 mM HEPES buffer, pH 7.4 in presence of 100mM NaCl and temperature 37°C, unless explicitly noted otherwise.

### ApoSOD1^2SH^ undergoes liquid-liquid phase separation

Using phase contrast microscopy (Fig. 2b-c, S2a), we did not observe the formation of condensates for different concentrations of WTSOD1^2SH^ (25, 50, 100, and 200 μM of WTSOD1^2SH^). We also tried to induce LLPS by adding heparin - a polyanionic glycosaminoglycan known to be an aggregation inducing agent (23, 24), which has also been shown to drive phase separation of proteins with low complexity regions (LCRs) or IDRs like Tau (25), TDP43 LCD (26) and SH_3_-PRM_5_ (27). Droplet formation was not observed even in the presence of heparin for WTSOD1^2SH^ (Fig. S2a).

**Figure 2.**
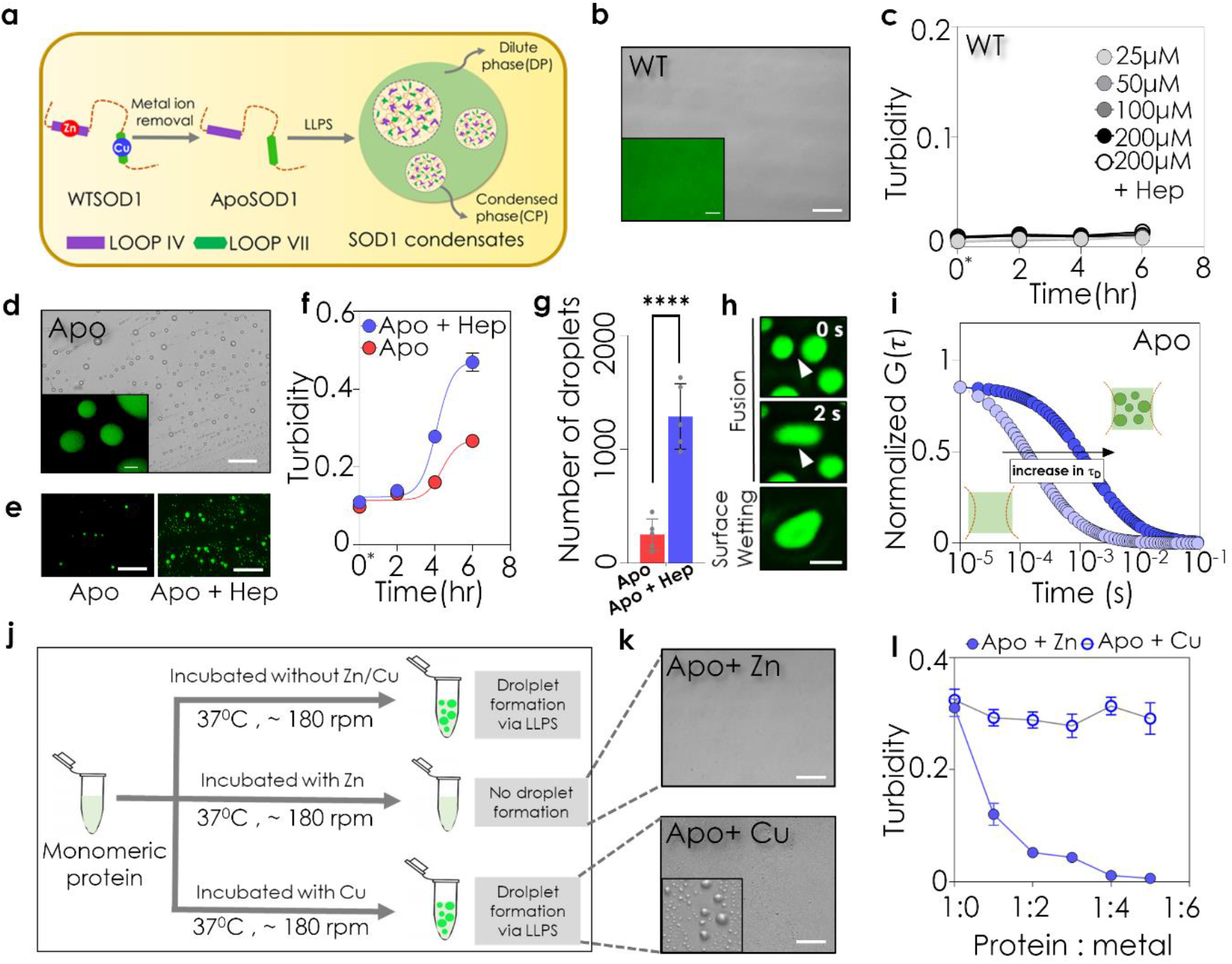
ApoSOD1^2SH^ undergoes liquid-liquid phase separation. **a)** SOD1^2SH^ undergoes LLPS on the removal of metal ion cofactors. **b)** DIC microscopic image of WTSOD1^2SH^ at 100 μM shows no droplet formation; scale bar: 100 μm inset shows Alexa Fluor 488 maleimide labelled WTSOD1^2SH^; scale bar: 5 μm **c)** Solution turbidity plot (absorbance at 600 nm) shows that WTSOD1^2SH^ (at concentrations ranging from 25 to 200 μM does not undergo LLPS in presence or absence of heparin, LLPS inducer. **d)** DIC microscopic image of ApoSOD1^2SH^ under condensate inducing conditions (37 °C, 180 RPM) at a concentration of 100 μM incubated in presence of 7% heparin and 100 mM NaCl; scale bar:100 μm (inset on bottom left shows Alexa Fluor 488 maleimide labeled protein condensates; scale bar: 5 μm. **e)** Fluorescent microscopic images of Alexa Fluor 488 maleimide labeled ApoSOD1^2SH^ condensates incubated in absence (left) and presence (right) of heparin at 2 hr timepoint; scale bar: 100 μm. **f)** Solution turbidity measurements with ApoSOD1^2SH^ (absorbance at 600 nm) shows that while in the absence of heparin the turbidity increases with time, the presence of heparin results in faster LLPS. **g)** Comparison of ApoSOD1^2SH^ droplet numbers calculated from 5 different images of droplets incubated with and without heparin at 2 hr timepoint. Statistical significance was established using unpaired Student’s t-test (****p< 0.0001). **h)** Liquid nature of ApoSOD1^2SH^ showing droplet fusion and surface wetting; scale bar: 5 μm **i)** Amplitude normalized FCS curves of ApoSOD1^2SH^ (in blue) of dilute phase (DP) and condensed phase (CP). Intense color indicates CP and light variant indicates DP. **j)** Scheme depicting that presence of Zn and not Cu in pre-incubation mixture inhibits LLPS. **k)** DIC microscopic images of ApoSOD1^2SH^ in presence of Zn (top) showing no droplet formation and Cu (below) showing the presence of droplets; scale bar: 100 μm. **j)** Solution turbidity plot (absorbance at 600 nm) of ApoSOD1^2SH^ subjected to increasing concentrations of Cu and Zn; turbidity decreases in a dose dependent manner for Zn while no significant change is observed in presence of Cu.

As it is known that immature SOD12SH exhibits conformational disorder in loop IV/VII that could promote LLPS, we next chose the metal-free Apo variant (ApoSOD1^2SH^) for further investigation. We removed both metal cofactors (Cu and Zn) by treating WTSOD1^2SH^ protein with EDTA using a previously described method (28). Interestingly, ApoSOD1^2SH^ formed spherical droplets in HEPES buffer at 37°C at concentrations greater than 2 μM (Fig. 2d). Fluorescence imaging of Alexa-488 Maleimide-labelled protein (20 nM labelled protein in the presence of 100 μM of unlabeled protein) showed that the protein molecules were distributed throughout the liquid droplets (Fig. 2d inset). We also find that the phase separation process was accelerated in the presence of heparin (Fig. 2e-g, S2b). With time, the droplets in proximity underwent fusion events forming larger droplets and surface wetting that could be monitored using fluorescence microscopy, exhibiting the liquid-like nature of these SOD1 condensates (Fig. 2h), like other proteins undergoing LLPS (29–33).

We used Fluorescence Correlation Spectroscopy (FCS) to monitor the change in the diffusional dynamics of the protein between the outside (dilute phase) and inside (condensed phase) droplets. FCS is a correlation analysis of temporal fluctuations of the fluorescence intensity at single molecule level in an approximately femtoliter confocal volume. From the value of the diffusion time (τ_D_), we calculated the hydrodynamic radius of ApoSOD1^2SH^ outside the droplet to be around 2.80 nm, which matched with earlier reports (34, 35). Inside the droplets, we observed a large (~2.5 fold) increase in the translational diffusion time (Fig. 2i). Interestingly, between the outside and inside droplets, we also observed a larger 17 times increase in the value of rotational correlation times (Fig. S2c), which we measured by time-resolved anisotropy measurements using Time Domain-Fluorescence Lifetime Imaging Microscopy (TD-FLIM). The restricted environment inside the droplet could be the reason for the increase in rotational as well as translational diffusion time of the protein molecules inside the droplets. It may be noted that a large decrease in diffusion coefficient between the solution phase and droplet has been previously observed for different proteins including FUS (36), TDP-43 (37), and Tau (38). In addition, we measured the fluorescence lifetime values of the attached dye (Alexa 488 Maleimide) to probe its excited state properties, which increased from dilute phase to condensed phase (Fig. S2d). When the condensates were subjected to 1,6-hexanediol treatment, an aliphatic alcohol that has been proposed to disrupt weak hydrophobic interactions (39), there was a significant reduction in the number of droplets (Fig. S2e-f). This implied the involvement of weak and non-specific hydrophobic interactions in stabilizing the ApoSOD1^2SH^ protein condensates.

### Absence of metal cofactor Zn (not Cu) is necessary for inducing LLPS

Since WTSOD1^2SH^ does not form condensates, while the metal-free ApoSOD1^2SH^ forms these readily, it is apparent that the metal cofactors may have a role to play in condensate formation. Since SOD1 contains two metals (Cu and Zn) as cofactors, we then wanted to test the effect of exogenous addition of each of these on the LLPS propensity of ApoSOD1^2SH^. We added Cu and Zn respectively to 100 μM protein solutions, which were then incubated for LLPS (pre-incubation condition, fig. 2j). We found that the addition of Zn at the time of incubation inhibited LLPS formation (Fig. 2j-l), while exogenous Cu addition did not have any effect on the condensate state of ApoSOD1^2SH^ (Fig. 2k-l). We then added Zn to pre-formed condensates of ApoSOD1^2SH^ (post-incubation condition) to check if Zn can dissolve them (Fig. S2g). This experiment clearly shows that LLPS of SOD1^2SH^ is reversible through Zn binding and this metal coordination can either inhibit (pre-incubation condition) or disrupt (post-incubation) condensates.

### ALS mutants with compromised Zn-binding also undergo LLPS

After establishing that Zn has a major role in modulating LLPS of SOD1^2SH^, we selected three ALS disease mutants for further studies. Two of these mutants, I113TSOD1^2SH^ and G85RSOD1^2SH^ are known to have compromised Zn binding in their native conditions (40, 41). The aberrant Zn binding of I113TSOD1^2SH^ has been attributed to the mutational stress introduced at the dimer surface of the protein (42). In G85RSOD1^2SH^, the mutation site is proximal to the Zn binding domain (residue 63-83). G85RSOD1^2SH^ was found to bind ~5% Zn relative to WTSOD1^2SH^ when expressed with the same metal ion supplementation (40). In the third ALS mutant G37RSOD1^2SH^ the mutation site is comparatively distant from the Zn binding site (23.7 Å). Although it has an intact Zn-binding site, G37RSOD1^2SH^ shows Cu deficiency (43). We have shown previously that G37RSOD1^2SH^ and I113TSOD1^2SH^ behave similarly to the WTSOD1^2SH^ and ApoSOD1^2SH^ respectively in terms of their membrane binding and aggregation propensity (44). In addition, the G37RSOD1^2SH^ mutant possesses a less severe disease phenotype (average patient survival time after disease onset ~17 years) as compared to I113TSOD1^2SH^ and G85RSOD1^2SH^ mutations (average survival times of ~4.3 years and 6 years, respectively) (45).

We found that the two variants, I113TSOD1^2SH^ and G85RSOD1^2SH^ formed spherical droplets while G37RSOD1^2SH^ did not (Fig. 3a, S3a). We observed that the addition of Cu did not have any effect on the condensed phase in either I113TSOD1^2SH^ or G85RSOD1^2SH^ (Fig.S3d-e). In contrast, the addition of Zn inhibited LLPS in I113TSOD1^2SH^ in a dose dependent way (Fig.3a, S3e). Interestingly, with G85RSOD1^2SH^, the addition of Zn did not lead to complete inhibition of LLPS (Fig. 3a, S3d-e) when compared to that of I113TSOD1^2SH.^ Time resolved anisotropy and FCS measurements confirmed that phase separated condensates of I113TSOD1^2SH^ and G85RSOD1^2SH^ were composed of slow diffusing species (Fig. 3b). The fluorescence lifetime of the probe increased inside the condensates (Fig. S3b-c). When we removed the metal ions from G37RSOD1^2SH^ by dialyzing the protein in EDTA, we found that the metal-free G37RSOD1^2SH^ mutant also readily underwent LLPS further highlighting the central role of metal binding in this process (Fig. 3c). The addition of Zn into metal-free G37RSOD1^2SH^ inhibited LLPS (Fig. 3c), while the addition of Cu did not yield any change (Fig. S3f). When Zn was added to pre-formed condensates (post-incubation condition) to see if the LLPS can be reversed (Fig. S3g), I113TSOD1^2SH^ droplets disappear completely whereas G85RSOD1^2SH^ droplets show partial dissolution (Fig. S3g). This experiment suggests that the LLPS of SOD1^2SH^ is effectively reversible through Zn coordination for ALS mutants as well unless the Zn binding is significantly compromised as in the case of G85RSOD1^2SH^.

**Figure 3.**
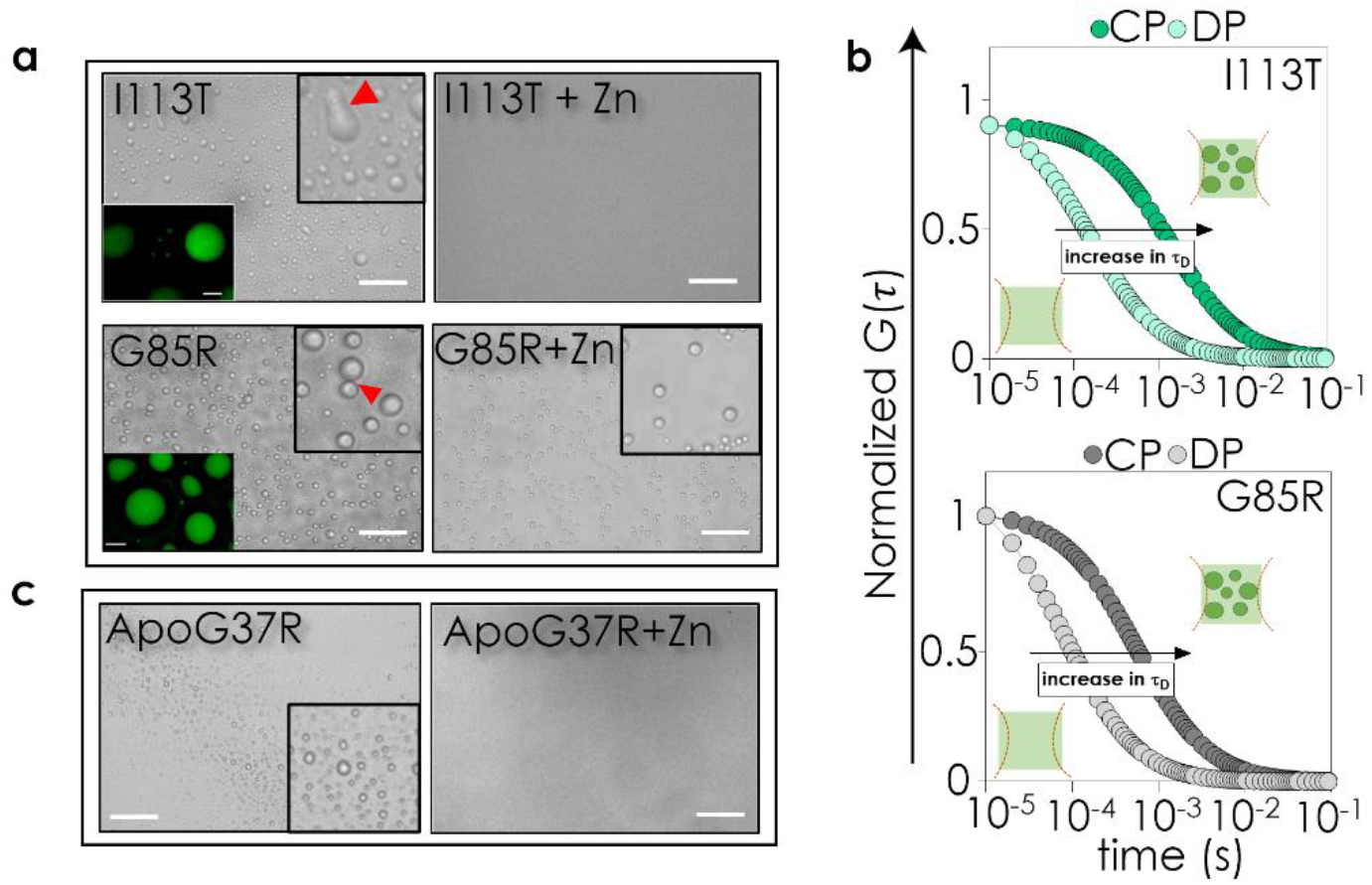
ALS mutants with compromised Zn binding undergo LLPS. **a)** DIC microscopic images of II13TSOD1^2SH^ droplets incubated in the absence (top left) and presence of Zn (top right) and G85RSOD1^2SH^ droplets incubated in the absence (bottom left) and presence (bottom right) of Zn respectively (scale bar: 100 μm); insets on bottom left show Alexa Fluor 488 maleimide labeled protein condensates for I113TSOD1^2SH^ and G85RSOD1^2SH^ respectively; scale bar: 5 μm. **b)** Amplitude normalized FCS curves of I113TSOD1^2SH^ (in green) and G85RSOD1^2SH^ (in grey) of dilute phase (DP) and condensed phase (CP) showing an increase in diffusion time from DP to CP. The intense color indicates CP and light variant indicates DP. **c)** On metal cofactor removal, ApoG37RSOD1^2SH^ formed condensates which ceased to exist when protein was incubated with Zn (left to right); scale bar: 100 μm.

### LLPS of monomeric SOD1 is driven by an order-to-disorder transition in the absence of Zn

The results above lead to a natural question about the mechanism of LLPS in the absence of Zn or reduced Zn-binding in the case of ALS mutants. NMR studies indicate that the conformational landscape of immature SOD1 comprises of a native-like state existing in equilibrium with four sparsely populated conformers (46), two of which can form non-native oligomers through the dimer interface and the electrostatic loop (loop - VII). It is possible that absence of Zn or reduced Zn-binding may give rise to such non-native interactions between immature SOD1 species and promote the formation of a liquid-like, condensed phase.

We used FCS to study the hydrodynamic radii of the different SOD1^2SH^ variants. Both ApoSOD1^2SH^ and severe mutants (I113TSOD1^2SH^ and G85RSOD1^2SH^) showed larger rH when compared to G37RSOD1^2SH^ and WTSOD1^2SH^ (Supplementary table 1). The value of rH obtained by FCS (this paper) for ApoSOD1^2SH^ matched well with previous experimental data (34, 35). To obtain further insights into this, we plotted the literature values (47) of rH of different proteins in their compact folded and chemically unfolded states. The log-log plot of the rH values of the folded proteins as a function of the number of residues could be fit to a straight line with a slope of 0.29 (Fig. 4b). For chemically unfolded proteins, the slope of the log-log plot was found to be 0.59. Interestingly, when we plotted the r_H_ values of these SOD1^2SH^ variants in Fig. 4b, we found that ApoSOD1^2SH^, I113TSOD1^2SH^ and G85RSOD1^2SH^ were positioned near the line for the chemically unfolded proteins. When we plotted the r_H_ value of α-synuclein, a well-characterized intrinsically disordered protein, it was found positioned near the same line. WTSOD1^2SH^ and G37RSOD1^2SH^ variants, on the other hand, were found near the line obtained for the compact folded proteins. This suggests that ApoSOD1^2SH^ and severe variants (I113TSOD1^2SH^ and G85RSOD1^2SH^) are more extended (or sample an extended conformer more often) compared to G37RSOD1^2SH^ and WTSOD1^2SH^ protein. We observed that rH decreased substantially with the addition of Zn in ApoSOD1^2SH^ and severe variants I113TSOD1^2SH^ and G85RSOD1^2SH^ (Fig. 4b, S4a-c).

**Figure 4.**
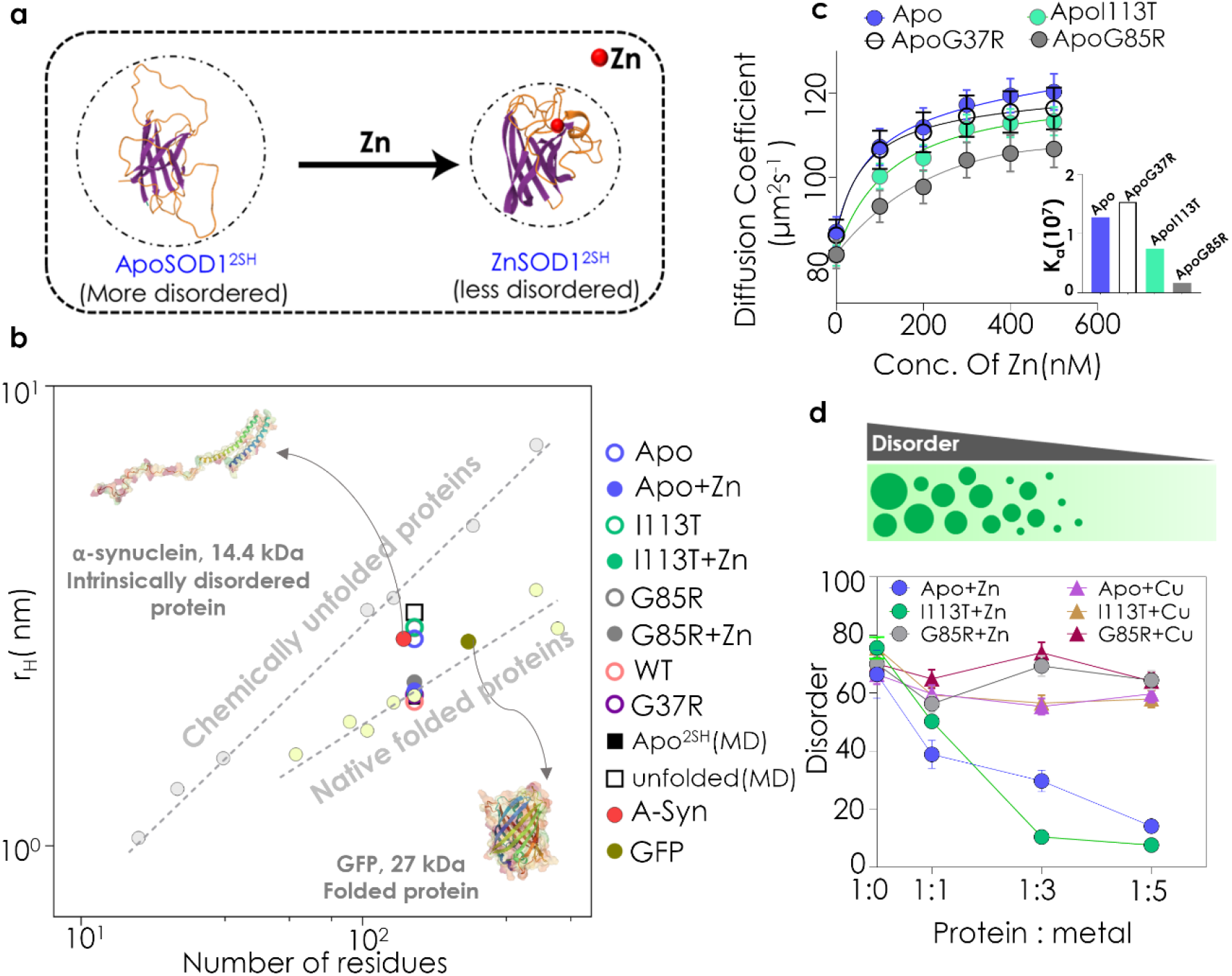
Zn drives disorder to order transition in SOD1. **a)** Scheme shows Zn dependent disorder to order transition in monomeric SOD1. **b)** Plot of the loge of the hydrodynamic radius (r_H_) versus the log e of the number of residues in the polypeptide chain. The line fitted to these data for the native folded proteins (yellow dashed) has a slope of 0.29 ± 0.02 and a y-axis intercept of 1.56 ± 0.1, while the other fitted to the chemically denatured protein (grey dashed) data has a slope of 0.57 ± 0.02 and a y-axis intercept of 0.79 ± 0.07. Literature data have been used for folded and chemically denatured proteins while we employed FCS to calculate the r_H_ of all SOD1^2SH^ variants with (solid shapes) and without Zn (hollow shapes). **c)** Variation of diffusion coefficients of de-metallated protein variants determined from FCS measurement with increasing Zn concentration (inset shows a comparison of binding constants between mutants and Zn; ApoSOD1^2SH^ and ApoG37RSOD1^2SH^ has a higher binding affinity to Zn than ApoI113TSOD1^2SH^ and ApoG85R SOD1^2SH^). **d)** Disordered/ extended conformation content decreases in ApoSOD1^2SH^ and ApoI113TSOD1^2SH^ with the addition of Zn calculated from FTIR spectra.

Since the addition of Zn resulted in large compaction in ApoSOD1^2SH^ and severe variants, we used FCS data to determine their Zn binding (Fig. 4c). For the Zn binding study, all protein variants were in their de-metallated Apo forms, which we prepared by dialysing the protein samples with EDTA using previous literature. We found that while ApoSOD1^2SH^ and ApoG37RSOD1^2SH^ bind to Zn with maximum affinity (k_a_ of 1.28×10^7^ M^-1^ and 1.54×10^7^ M^-1^ respectively) the binding of ApoG85RSOD1^2SH^ is the weakest (k_a_ of 1.71×10^6^ M^-1^). We then used FTIR to determine how the secondary structure of the protein changed as a result of cofactor insertion. Using FTIR, we found that disorder/extended conformation was higher in metal-free ApoSOD1^2SH^ and severe variants (I113TSOD1^2SH^ and G85RSOD1^2SH^) when compared to WTSOD1^2SH^ and G37RSOD1^2SH^ (Fig. S4d, Supplementary table 2). The disordered/extended components in ApoSOD1^2SH^ and I113TSOD1^2SH^ (Fig. 4d, S4e-f) decreased as Zn was added, while the effect of Zn was less in G85RSOD1^2SH^ (Fig. S4g). No significant change in the secondary structure for any variant was noted in presence of Cu (Fig. 4d, S4e-g). Together, FTIR and FCS data revealed that the addition of Zn in ApoSOD1^2SH^ and I113SOD1^2SH^ resulted in a conformational transition of the Zn free protein from an extended (FCS data, greater r_H_) disordered (FT-IR data) state to a compact (lower r_H_) and less disordered state (FT-IR data). For G85RSOD1^2SH^, this transition is somewhat partial presumably because of its weaker zinc binding compared to the I113T mutant.

### Characterization of disorder in monomeric SOD1 and condensed phase interactions using simulations

To complement FITR and FCS experiments, we characterized the extent of conformational disorder within ensembles of SOD1 monomer variants obtained from atomistic MD simulations. Simulations were performed using the AMBER99SB-disp force field which was shown to be suitable for simulating both folded and disordered proteins (see Methods). The hydrodynamic radius r_H_ was calculated using the HullRad algorithm (48) which employs a convex hull method to estimate the hydrodynamic volume of the protein molecule. The mean r_H_ value ranges from 2.1 nm for ApoSOD1^2SH^ variants to 3.2 nm for unfolded SOD1^2SH^ (Fig 4b), which suggests that the extent of structural disorder observed in FCS measurements for ApoSOD1^2SH^ variants is intermediate to those of the simulated monomeric states. Secondary structure analysis of ApoSOD1^2SH^ variants using the DSSP algorithm (49) indicates the presence of coil-like conformations in loop-IV/VII while β-sheet conformations were dominant in the expected β-barrel regions (Fig. S5a). In contrast, unfolded SOD1^2SH^ largely adopts a coil conformation with short stretches of α-helix and β-sheet conformations (Fig. S5b). Distance root mean square deviation (dRMSD) analysis confirms that the β-barrel fold is stable (dRMSD <=0.3 nm) across all ApoSOD1^2SH^ variants (Fig. S5c) which is consistent with previous NMR/SAXS experiments (50). Representative conformations of ApoSOD1^2SH^ and unfolded SOD1^2SH^ are shown in Fig. S5d.

Consistent with secondary structure and dRMSD analysis, root mean square fluctuation (RMSF) of Cα atoms suggests that ApoSOD1^2SH^ variants exhibit high conformational flexibility (RMSF_max_> 0.4 nm) in loop-IV/VII regions (Fig. 5a). To assess the effect of Zn binding on conformational disorder of monomeric SOD1, we also simulated three additional immature states: ApoSOD1^S-S^ (disulfide present, without metal cofactors), ZnSOD1^2SH^ (Zn-bound and disulfide reduced) and ZnSOD1^S-S^ (Zn-bound and disulfide present). We compared the conformational flexibility across different regions of the SOD1 by analyzing their RMSF profiles (Fig. 5b). The RMSF profile of ApoSOD1^2SH^ shows considerable flexibility in loop-IV, and loop-VII wherein RMSF for several residues exceed 0.3 nm. Notably, the presence of disulfide bond only reduced flexibility in the aa:51-60 region (RMSF_max_~0.15 nm) within loop-IV which is involved in disulfide bond formation through C57 while loop-VII remained flexible. Zn abolished the flexibility of loop-IV (RMSF_max_~0.15 nm) to which it directly binds and led to a reduction in the RMSF of loop-VII residues. In conclusion, the high flexibility of loop-IV/VII observed in the absence of Zn in simulations is consistent with experimental observations and points towards an intrinsic propensity for structural disorder within the SOD1 monomer in the absence of Zn.

**Figure 5.**
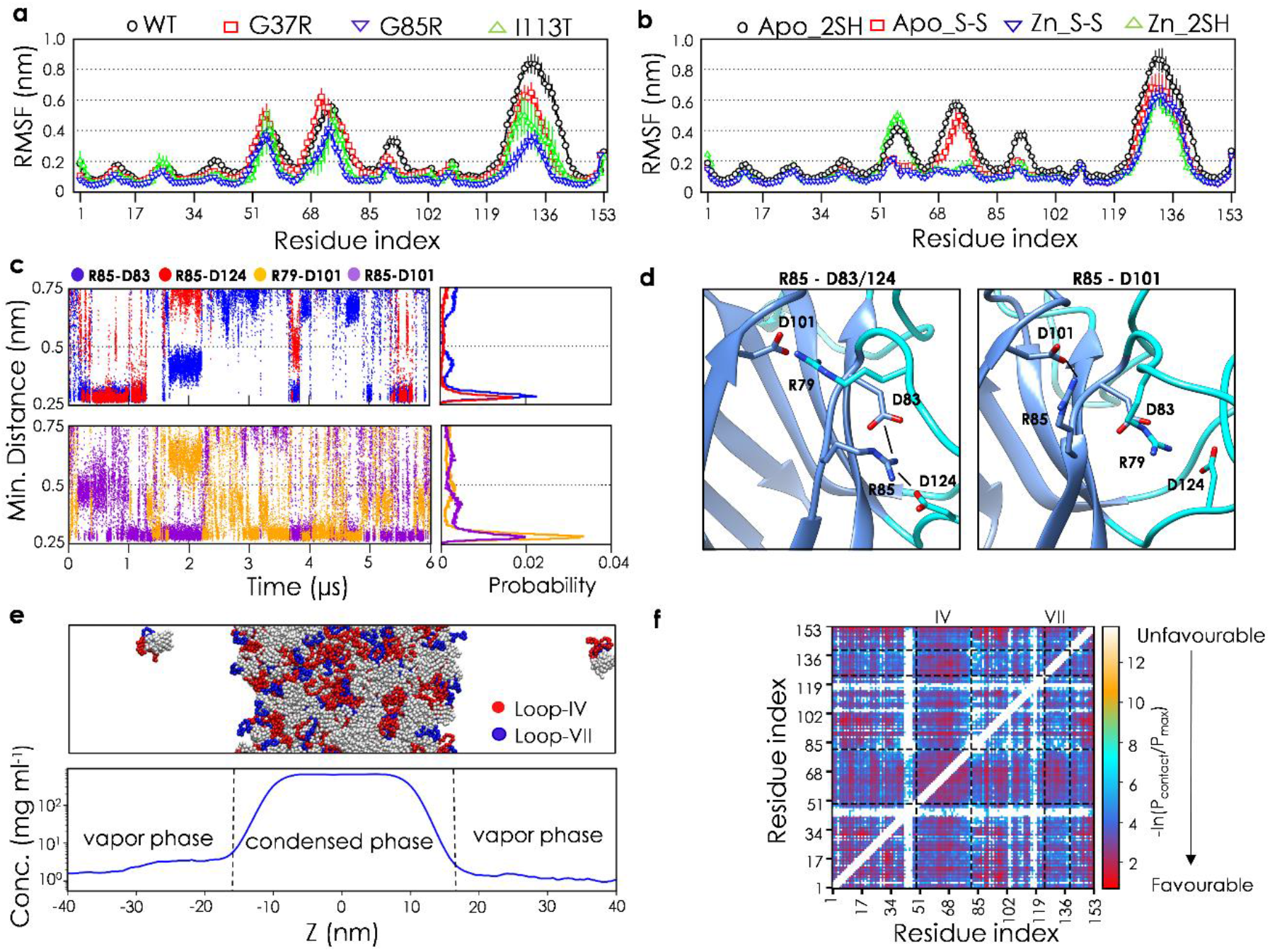
Computational characterization of SOD1^2SH^ variants and molecular interactions in the condensed phase. **a)** Per-residue RMSF profiles of ApoSOD1^2SH^, WTSOD1^2SH^ & mutants. **b)** Per-residue RMSF profiles of Apo SOD1^2SH^, SOD1^S-S^, ZnSOD1^S-S^ and ZnSOD1^2SH^. **c)** Salt-bridge analysis of G85RSOD1 Apo^2SH^ (dist. cutoff <0.5 nm = salt-bridge). **d)** Snapshots from the G85RSOD1 Apo^2SH^ trajectory showing the formation of non-native salt-bridges which are likely to be detrimental for Zn-binding. **e)** (top) Slab configuration for CG simulations of partial rigid SOD1 at 275K (red circle represents the loop IV, blue circle represents loop VII, green color represents the flexible domains. The condensed phase is centered in the middle of the simulation box. (bottom) Density profile of the CG slab simulation. **f)** Free energy profile of pairwise intermolecular contact formation (top) and total contact probabilities per residue (bottom) in the condensed phase as a function of residue index.

To understand the mechanism by which G85RSOD1^2SH^ substitution renders SOD1^2SH^ incapable of binding to Zn, we analyzed the interactions of R85 with neighboring residues over the course of the 6 μs run (Fig. 5c & 5d). We observed that R85 formed salt-bridges with key aspartate residues (51) which either stabilize Zn (D83/D124) or may play a role in stabilizing the native conformation of loop-IV (D101). D83 directly binds to Zn whereas D124 coordinates two key histidine residues involved in Zn coordination. D101 natively forms a salt-bridge with R79 which may help stabilize the conformation of loop-IV. Overall, simulation of ApoG85RSOD1^2SH^ suggests a mechanism wherein the formation of dynamic, non-native salt-bridges involving R85 may hinder its ability to bind Zn. Induction of non-native, intramolecular salt-bridges through post-translational modifications (52–54) have been previously shown to alter the binding affinity of protein-protein interactions and may therefore also modulate ApoSOD12^SH^Zn^2+^ affinity through a similar mechanism via G85R mutation.

In order to determine which regions ApoSOD1^2SH^ are involved in stabilizing the condensate, we performed a coarse-grained (CG), phase coexistence simulation (Fig. 5e, top) using a protocol which has been described previously (55). Briefly, ApoSOD1^2SH^ monomers were modelled at a single bead per amino acid resolution and formed a stable condensate at 275 K with the CG model (Fig. 5e, bottom). To model ApoSOD1^2SH^, the residues which form the β-barrel structure were made rigid relative to each other while those belonging to loop -IV/VII were allowed to remain flexible. The analysis of intermolecular contact maps between SOD1 monomers in the condensed phase indicate that several regions of the β-barrel and disordered loop-IV (aa:60-75) collectively stabilize the condensate. In contrast, significantly fewer intermolecular contacts were observed for loop-VII (Fig. 5f, Supplementary movie 1). The classification of the observed contacts based on residue type indicate predominance of charged interactions between lysine and aspartate/glutamate residues (Fig. S5e & S5f). Among the hydrophobic residues, valine/isoleucine interacted with lysine. Polar interactions between serine-glycine and serine-lysine pairs were also observed. Taken together, the CG condensate simulation suggests that ApoSOD1^2SH^ condensate is predominantly stabilized through a combination of both electrostatic and hydrophobic interactions involving the β-barrel and loop-IV.

### LLPS of SOD1 is followed by aggregation

Studies of RNA-binding proteins (56–58) involved in SG formation such as FUS, hnRNPA1/2 and TDP-43 indicate that LLPS of their disordered, low complexity domains (LCDs) gives rise to labile fibrils upon droplet maturation. Moreover, the liquid-to-solid phase transition which leads to the formation of fibrils was shown to be enhanced by mutations associated with neurodegenerative diseases. Based on these observations, it has been proposed that LLPS may represent an intermediate state en route to the toxic aggregate formation (59).

After establishing that ApoSOD1^2SH^ and severe ALS mutants (I113TSOD1^2SH^ and G85RSOD1^2SH^) undergo a Zn-dependent phase separation, we investigated the temporal maturation of ApoSOD1^2SH^ droplets using fluorescence microscopy in combination with FCS (Fig. 6a-b). As the maturation progressed with time, we observed the presence of high intensity fluorescent aggregates which started forming from within the droplets (Fig. 6c). FCS experiments were carried out at different regions and different time points to investigate the diffusional dynamics of droplets maturation. For the FCS experiments, we chose three regions, namely the diffused part of the image outside the droplets (region 1, Fig 6a-c, S6a); the relatively low intensity portion inside the droplets (region 2, Fig. 6a-c, S6b); and the high intensity regions inside the droplets, which contained the aggregates (region 3, Fig. 6a-c). The analyses of the correlation functions obtained at region 1 showed the presence of fast diffusing molecules (Fig. 6d). The hydrodynamic properties (r_H_ of 2.80 nm) of these fast-moving molecules resembled that of labeled ApoSOD1^2SH^, suggesting that region 1 predominantly contained monomeric protein.

**Figure 6.**
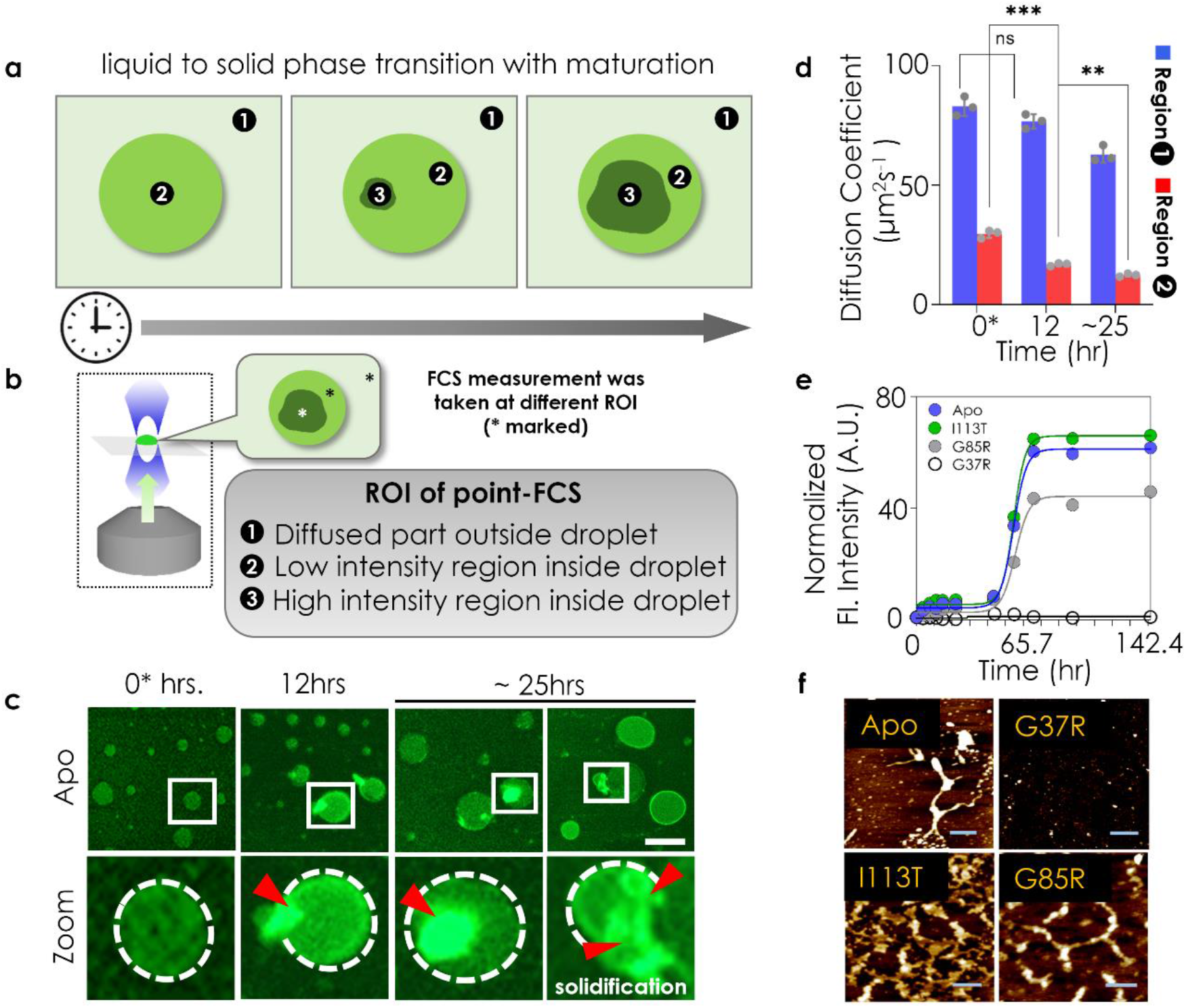
Maturation of liquid droplets precedes aggregation. **a)** Cartoon scheme showing the maturation of protein droplet with time (region1: diffused region outside droplet, region 2: low intensity portion inside the droplet, region 3: highly intense portion inside droplet appears during maturation) at different time points. **b)** Diffusion coefficient of region 1 and 2 (within droplet) was measured using point FCS as shown in the scheme. **c)** Upper panel shows fluorescence images of the maturation of Alexa Fluor 488 labeled ApoSOD1^2SH^ droplets with time, samples were incubated at 37 °C up to ~25 hours. Images are taken at indicated time point: 0* hour (the time after incubation required for droplet formation), 12 hours and >24 hours; images below show magnified scale bar: 5 μm. The lower panel shows magnified view of the marked region from corresponding images on the upper panel. **d)** Diffusion coefficient obtained from point FCS measurement at different regions of the droplet during maturation course; 0* (x-axis label) indicates the time after incubation required for droplet formation. Statistical significance was measured using paired Students t-test; ns denotes non-significant; **p< 0.01; ***p< 0.001 **e)** ThT fluorescence assay plot showing aggregation of all SOD1^2SH^ variants. ApoSOD1^2SH^, I113TSOD1^2SH^ and G85RSOD1^2SH^ show high ThT fluorescence. **f)** AFM micrographs of protein variants show fibrillar aggregates in ApoSOD1^2SH^, I113TSOD1^2SH^ and G85RSOD1^2SH^ after prolonged incubation at 37°C for 144 hours; scale bar: 200 nm.

In contrast, analyses of the correlation functions obtained inside the droplets (region 2) were complex. Even at the early time points (for example, at 0* hrs in fig. 6d, which shows the data at the earliest time point when LLPS was observed) the values of D were found considerably less than what was observed for the monomeric protein in solution or for the molecules present at the diffused region (region 1) indicating restricting environment inside the droplets. As the maturation progresses, the values of D decreased further with time (for example 12 hours in Fig. 6d). Interestingly, at 24 hours and beyond, the correlation functions contained more than one component with a large fraction of a very low diffusing component (Fig. 6d) The region containing the aggregates (region 3) did not yield any correlation functions as their diffusion behavior was too slow to measure.

### SOD1 aggregates are toxic and not reversible by Zn addition

We then used ThT fluorescence assay to determine if the aggregates obtained after long incubation would eventually lead the to amyloid formation or these are amorphous. We found that the ApoSOD1^2SH^ and two severe mutants (I113TSOD1^2SH^ and G85RSOD1^2SH^) showed large ThT fluorescence indicating amyloid formation, while G37RSOD1^2SH^ did not aggregate (Fig. 6e). The aggregation kinetics were sigmoidal with prominent lag phases. When we added a small concentration (10 μM) of pre-aggregated ApoSOD1^2SH^ at the beginning of the aggregation assay, the kinetics became hyperbolic (Fig. S6c), which is typical of seeded aggregation. Interestingly, when we added the same concentration of ApoSOD1^2SH^ condensate as the seed at the beginning, the kinetics remained sigmoidal indicating that the condensates were not able to seed amyloid formation (Fig S6c). We then used AFM experiments to complement ThT fluorescence data (Fig. 6f, S6d), which provided information about the height and morphology of the aggregates. AFM micrographs of ApoSOD1^2SH^, I113TSOD1^2SH^ and G85RSOD1^2SH^ aggregates showed the presence of long, dense fibrils (Fig. 6f, S6d).

We then wanted to find out if the protein variants induced toxicity in their condensate and aggregate states. We first used a membrane permeation assay in which we used calcein encapsulated SUVs and monitored the degree of calcein leakage upon addition of the proteins. In this assay, the protein variants were subjected to the solution conditions to induce LLPS formation and aggregation. Upon confirmation using microscopy, the protein samples in their respective conditions were studied for their pore forming propensities. We found that ApoSOD1^2SH^, I113TSOD1^2SH^ and G85RSOD1^2SH^ aggregates induced significant vesicle damage when compared to their condensate forms (Fig. S6e). The calcein release assay-which provides an estimate of the membrane permeability-was then complemented by measuring cellular toxicity using MTT assay in SHSY5Y neuroblastoma cells (Fig. S6f). Again, I113TSOD1^2SH^ and G85RSOD1^2SH^ aggregates were found to be detrimental to cell viability, while the condensates were found to be less toxic (Fig. S6f). However, it has also been reported that the mutants of prion-like (PRD) domain of TDP-43 that are aggregation prone confer less toxicity to yeast cells than mutants that promote condensate formation (60).

We had observed earlier that LLPS was completely reversible when Zn was added either at the pre-incubation or post-incubation conditions (Fig. 1). Therefore, we chose to probe the effect of the metal cofactors on the formation and stability of amyloid aggregates under pre-incubation and post-incubation conditions respectively. When we added Zn at the beginning of incubation (pre-incubation), aggregation of ApoSOD1^2SH^ and I113TSOD1^2SH^ (Fig. S6g-h) was inhibited while the presence of Cu did not suppress aggregation (Fig. S6i-j). Interestingly, short fibrils were observed for G85RSOD1^2SH^ incubated with Zn (Fig. S6h) which is consistent with its significantly reduced affinity towards Zn. In contrast, when we added Zn at the stationary phase of the amyloid formation (144 hours, post-incubation), we found no significant change either in ThT fluorescence (Fig. S6k) or AFM (Fig. S6l). These data in combination with Zn-induced dissolution of ApoSOD1^2SH^ droplets clearly suggest that while the state of droplet formation is reversible by Zn binding, the protein aggregates are stable and not altered by either of the cofactors.

## Discussion

Using both experiments and molecular simulations, we show here that while compact, WTSOD1^2SH^ does not undergo phase separation, partially disordered ApoSOD1^2SH^ or Zn-compromised mutants engage in non-native intermolecular interactions which lead to condensate formation. Unlike ApoSOD1^2SH^ and I113TSOD1^2SH^, G85RSOD1^2SH^ exhibits a significant reduction in Zn binding affinity due to non-native salt-bridge formation as substantiated by FCS and atomistic simulation data. A coarse-grained simulation of the condensed phase suggests that the droplets are predominantly stabilized by residues of β barrel and loop-IV through electrostatic and hydrophobic interactions. Prolonged incubation of the condensates facilitates a liquid to solid-like transition which culminates in aggregation. While these aggregates are toxic and stable, condensates do not exhibit significant toxicity and can be dissolved through the addition of Zn.

Our model of ‘cofactor regulated LLPS of SOD1’ proposes that Zn (and not Cu) removal induces sampling of disordered states, LLPS, and favorable conversion to amyloids (Fig. 7). Our results also correlate with the severity in disease phenotypes for two ALS disease mutants under in vitro conditions. The less severe G37RSOD1^2SH^ mutant is compact with a lower tendency towards LLPS. In contrast, the more severe I113TSOD1^2SH^ and G85RSOD1^2SH^ mutants, with reduced Zn affinity are more disordered, readily forming droplets and amyloids. Overall, our results highlight that Zn is a key regulator of the LLPS process, which shields the loop regions and suppresses the aberrant accumulation of conformationally labile molecules, thereby preventing LLPS.

**Figure 7.**
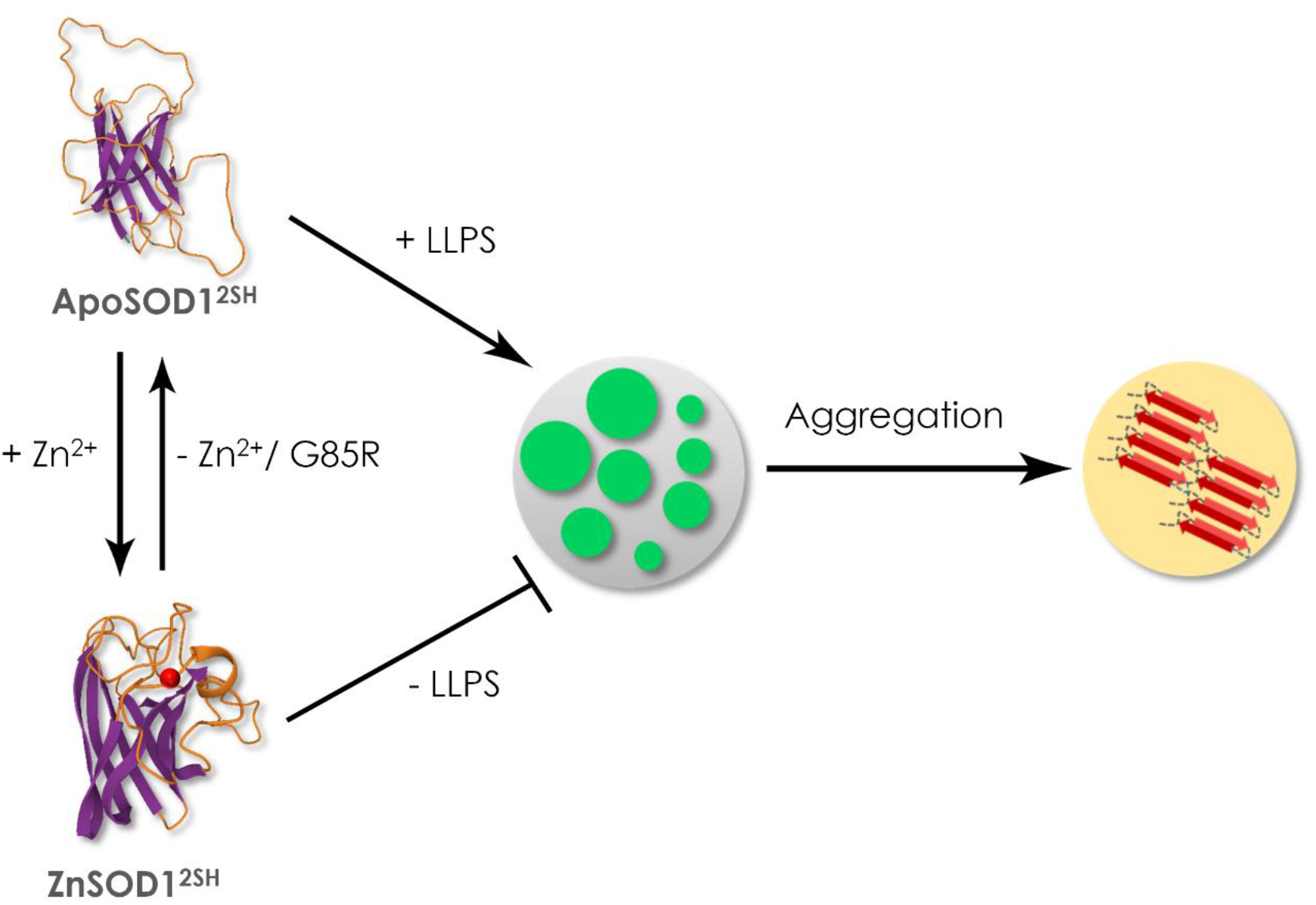
Schematic showing a model of Zn regulated LLPS leading to aggregation.

It is important to consider that intracellular aggregates of mutant SOD1 and aggregates isolated from spinal cord tissue of ALS-SOD1 mouse models were found to be metal deficient (61). The ability to investigate metal-binding in mutant aggregates from patient samples is limited since fALS only accounts for 5-10% of ALS cases, while most instances of the disease are sporadic (90-95%) (62). Sporadic ALS may possibly stem from toxic intermediates of the reduced Apo form due to failure in metal uptake or oxidation. We believe that cofactor regulated LLPS of SOD1 will play an important role in SOD1 aggregation pathology and ALS disease biology. These findings accentuate the need to delve deeper into the cellular events that trigger disrupted or improper metal loading to nascent SOD1 resulting in compromised stability and formation of non-native intermediates that switch to the aggregation pathway to enhance our understanding of ALS disease biology.

Kay and colleagues reported that post-translational modifications during protein maturation such as metallation and oxidation assist in eliminating off-pathway conformers that correspond to high energy states in immature SOD1 (63). From an evolutionary perspective, it might be tempting to speculate whether these off-pathway intermediates are sequestered into condensates as a transient neuroprotective mechanism against aggregation. In direct contradiction with the idea of toxic aggregates, however, are observations from recent studies (60,33) on TDP-43 C-terminal domain and Tau which indicate that condensates promote cellular toxicity as opposed to aggregates. Further studies are needed to determine the relative significance of condensate versus aggregate toxicity for proteins implicated in neurodegenerative diseases.

As discussed earlier, SOD1 SGs and inclusion bodies are found in the spinal cords of ALS patients. Based on recent LLPS studies and the data presented here, we speculate that these may originate via condensate formation and compartmentalize toxic species. A detailed understanding of how condensates metamorphose to inclusion bodies could open possibilities for developing therapeutic drugs targeting these non-native intermediates instead of aggregate species.

## Materials and Methods

Refer to SI Appendix for details. Recombinant SOD12SH was purified in E. coli (BL21 DE3 strain). Cells were grown in Luria Bertani media and 1 M isopropyl-1-thio-β-d-galactopyranoside was used to induce overexpression of the protein. 1 mM CuSO4 was added for proper metal loading. Cells were harvested by centrifugation and re-suspended in pre-chilled lysis buffer (20 mM Tris–HCl, 500 mM NaCl, pH 8.0). After removal of cell debris by centrifugation, the supernatant was loaded onto Ni-NTA column. The column was washed using 50 ml wash buffer (20 mM Tris–HCl, 500 mM NaCl and 50 mM imidazole, pH 8.0) followed by elution with 20 mM Tris–HCl, 500 mM NaCl and 500 mM imidazole, pH 8.0. Eluted fractions were pooled and dialysed using SnakeSkin Dialysis Tubing (10 KDa MWCO) in 20 mM Na-phosphate buffer pH 7.4. The protein concentration was estimated by recording absorbance at 280 nm using the molar extinction coefficient (ε280= 5500 cm^-1^). ApoSOD12SH was prepared from the holoSOD12SH by metal removal using EDTA. All SOD12SH protein variants were labelled using a AlexaFluor 488 Protein Labelling kit (Invitrogen). Confocal microscopy, fluorescence correlation spectroscopy and Fourier-transform infrared spectroscopy were used to characterize phase behavior. TD-FLIM and FCS were used to study droplet maturation. Protein aggregation kinetics was studied via ThT assay and aggregate morphology was studied using atomic force microscopy. For calcein release assay, calcein-loaded SUVs composed of 3:7 POPC: DOPS were added to SOD12SH condensates and aggregates at a protein: lipid ratio of 1:10. 50 mM calcein was encapsulated within the SUVs. Samples were excited at 490 nm. 1 μl Triton X-100 was used to determine 100% calcein release, and all results were normalized to this value. For MTT assay, SHSY5Y neuroblastoma cell line was used. Cells were given a 12 hrs treatment with 5 μM protein condensates and aggregates, followed by a 4 hr incubation with MTT solution and absorbance measurement at 595 nm using an ELISA reader (Emax, Molecular Device). MD simulations were conducted to determine molecular level details of atomic interactions during phase separation.

## Acknowledgments

We thank T. Murugunandan and S. Laha for technical support in Atomic Force Microscopy and Fourier Transform Infrared Spectroscopy; Central Instrumentation Facility, Indian Institute of Chemical Biology for provision of infrastructure; Bishal Roy and Subhadeep Das for image-processing; work done at Indian Institute of Chemical Biology was supported by Council of Scientific & Industrial Research and University Grants Commission. Work done at Texas A&M University was supported by NINDS and NIA R01NS116176. All-atom simulations were performed on the Extreme Science and Engineering Discovery Environment (XSEDE), which is supported by National Science Foundation grant number ACI-1548562. The simulations utilised the XSEDE Expanse GPU at the San Diego Supercomputer Center through allocation TG-MCB120014.

