## Supplementary Information for "Zn-dependent structural transition of SOD1 modulates its ability to undergo liquid-liquid phase separation"

**This PDF file includes:**

S1 to S6

Supplementary tables 1 & 2

Supplementary movie 1

Material and Methods

SI References

### 1. Supplementary Figures

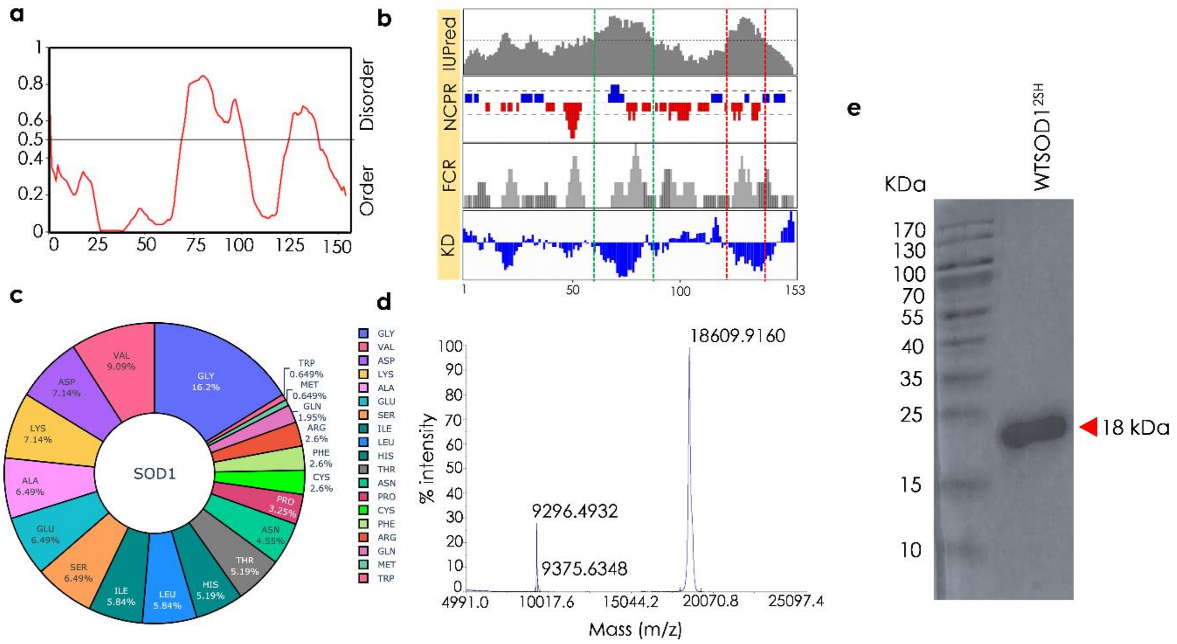


**Figure S1: a)** Disorder propensity of SOD1 residues generated by PONDR. SOD1 has two disordered stretches extending from 70 to 101 and 125 to 139. **b)** Linear sequence analysis of SOD1 (PDB ID: 4BCY) . Each track reports a different type of sequence feature to provide a general summary of the linear amino acid sequence. IUPred represents the predicted disorder score based on the IUPred algorithm; The dotted horizontal line indicates the threshold (0.5) for disorder. Amino acids sequence ranging from 63 to 83 (Zn binding loop) & loop 120-141 (the electrostatic loop) are identified to be intrinsically disordered (IDRs). NCPR profile song the linear sequence of SOD1 represents the linear net charge per residue; in which the Zn binding domain (ZBD) shows both net positively charged local regions and net negatively charged regions, suggesting electrostatic interactions may play a role in driving ZBD-ZBD interaction. The blue and red peaks denote positive and negative charges, respectively. FCR represents the fraction of charged residues, while KD represents the Kyte-Doolittle hydrophobicity scale. We found that the Zn binding and Electrostatic loops are significantly less hydrophobic. FCR, NCPR and KD plots were obtained by CIDER. **c)** Amino acid composition of SOD1 with respect to the hydrophobicity and charge. **d)** Matrix-assisted laser desorption ionization-time of flight (MALDI-TOF) mass spectrum of SOD1$^{2SH}$. The intense peak at m/z ~18,600 (Da) corresponds to the molecular mass of the intact monomeric protein. The less intense peak at m/z ~9,300 (Da) is half that of the molecular mass peak and is due to a doubly charged molecular ion. E) Denaturing SDS gel shows monomeric SOD1$^{2SH}$ band at ~18 kDa.

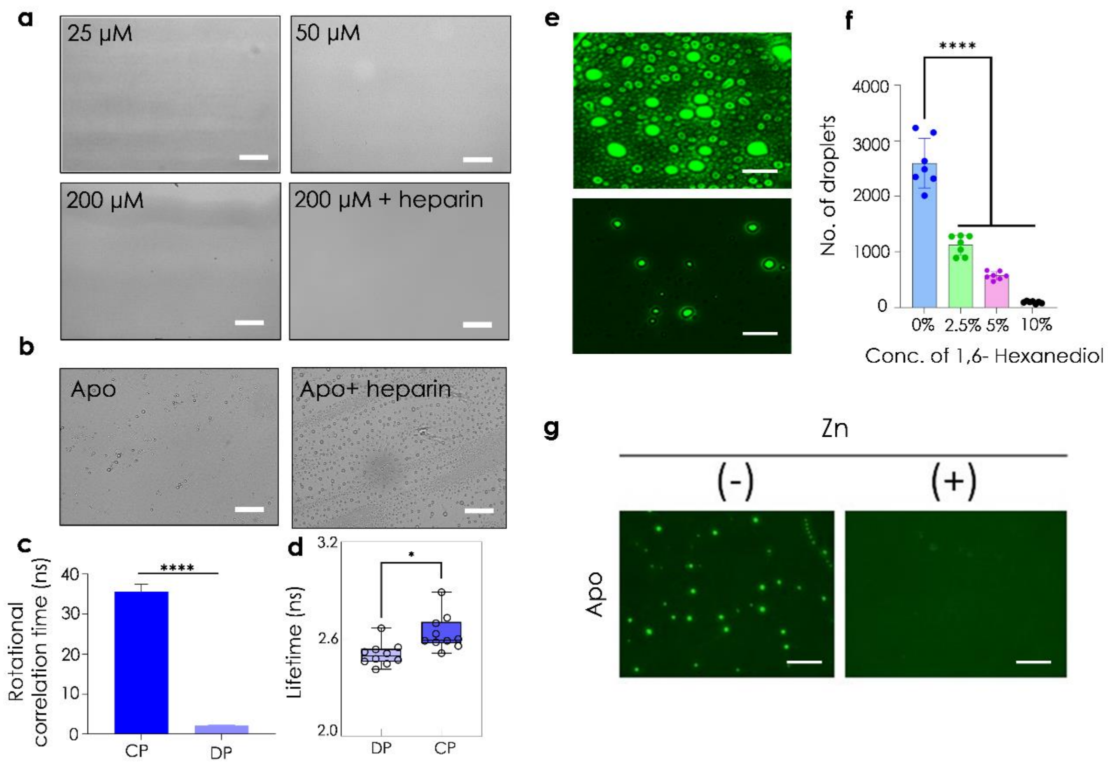


**Figure S2:** a) Figure panel shows DIC microscopic images of WTSOD1$^{2SH}$ (at concentrations 25, 50, 200 µM and 200 µM incubated with heparin) does not undergo liquid-liquid phase separation; scale bar: 100 µm. b) DIC images of ApoSOD1$^{2SH}$ droplets in the absence (left) and presence of heparin (right) at 2 hours timepoint; scale bar: 100 µm. c) Rotational correlation time values from time resolved anisotropy measurements of ApoSOD1$^{2SH}$ condensates (DP: dilute phase; CP: condensed phase); **** p< 0.0001. d) Fluorescence lifetime values for condensed (CP) and dilute phases (DP) of ApoSOD1$^{2SH}$; *p<0.05. e) Fluorescent images of liquid droplets of Alexa Fluor 488 maleimide labelled ApoSOD1$^{2SH}$ in absence (top) and presence (bottom) of 10% 1,6 –hexanediol; scale bar: 10 µm. f) Mean number of droplets calculated from 7 different DIC microscopic images of apoSOD1$^{2SH}$ in presence of hexanediol at mentioned concentrations; ****p<0.0001 g) fluorescent images of ApoSOD1$^{2SH}$ droplets in presence and absence of Zn show that addition of Zn to preformed droplets resulted in droplet dissolution in ApoSOD1$^{2SH}$; scale bar: 100 µm.

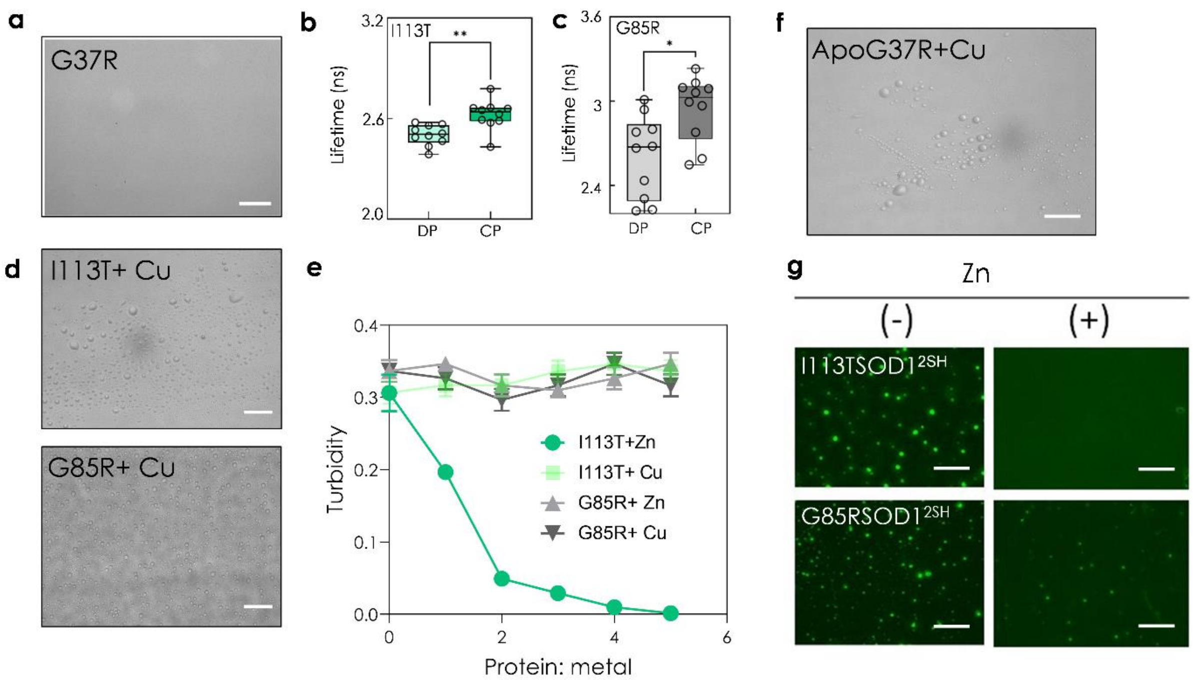


**Figure S3:** a) DIC microscopic image of G37RSOD1$^{2SH}$ showing no LLPS; scale bar: 100 µm. b) Fluorescence lifetime values for condensed and dilute phases of I113TSOD1$^{2SH}$ (in green); **p<0.01 and c) G85RSOD1$^{2SH}$ (in grey) *p<0.05 from TD-FLIM measurements; CP and DP indicate condensed and dilute phases respectively. d) DIC microscopic images of I113TSOD1$^{2SH}$ (top) and G85RSOD1$^{2SH}$ (below) droplets in presence of Cu; scale bar: 100 µm. e) Solution turbidity (absorbance at 600 nm) of I113TSOD1$^{2SH}$ and G85RSOD1$^{2SH}$ in presence of increasing concentrations of Zn and Cu. f) DIC image of ApoG37RSOD1$^{2SH}$ in presence of Cu showing LLPS; scale bar: 100 µm. g) fluorescent images of I113TSOD1$^{2SH}$ and G85RSOD1$^{2SH}$ droplets in absence and presence of Zn; scale bar: 100 µm.

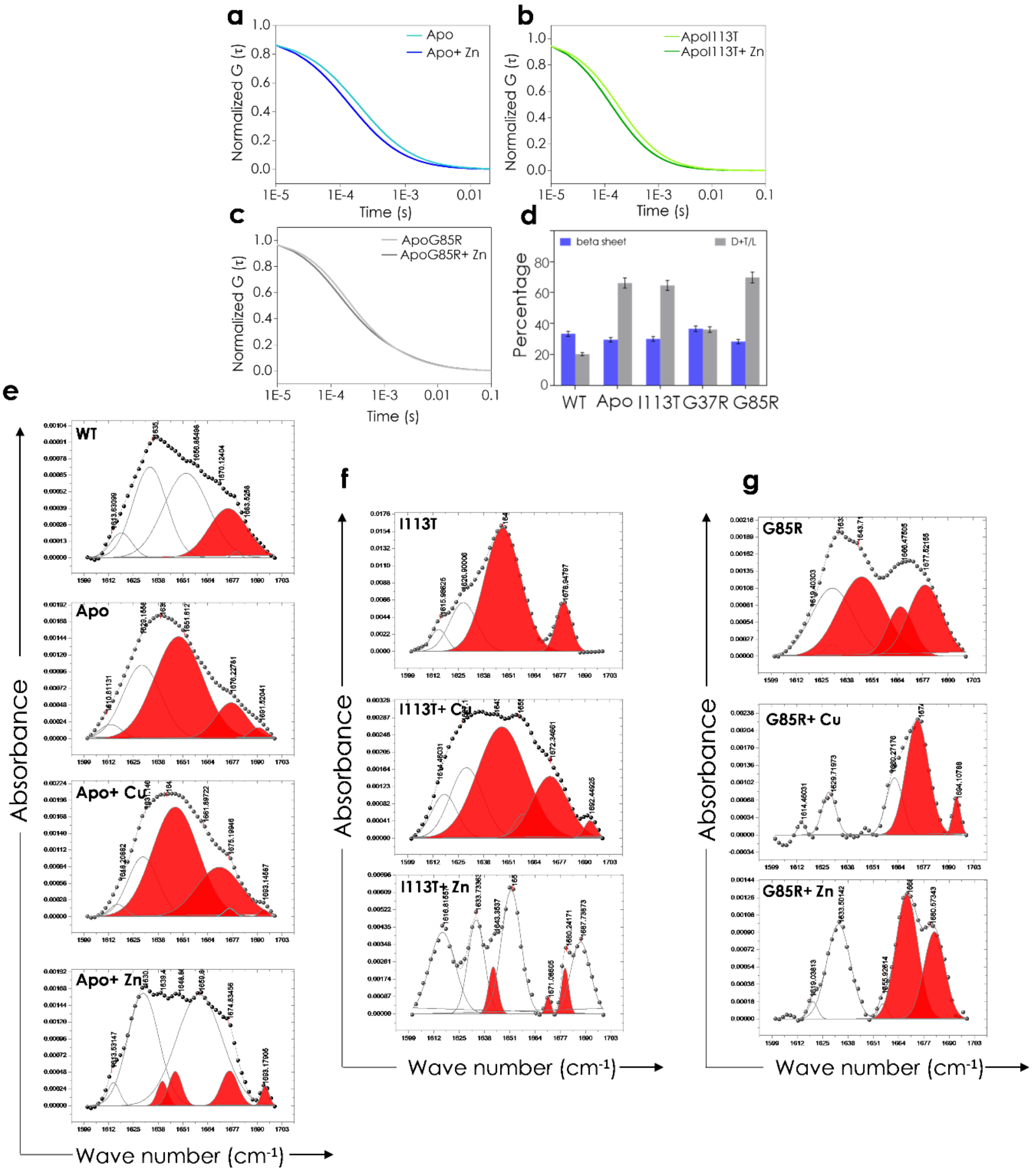


**Figure S4: Normalized FCS curves of** a) ApoSOD1$^{2SH}$ b) ApoI113TSOD1$^{2SH}$ and c) ApoG85RSOD1$^{2SH}$ in presence and absence of Zn. d) Comparison of beta sheet content and disorder/loops and turns among SOD1$^{2SH}$ mutants obtained from FTIR; ApoSOD1$^{2SH}$, I113TSOD1$^{2SH}$ and G85RSOD1$^{2SH}$ show higher disordered conformation than WTSOD1$^{2SH}$ and G37RSOD1$^{2SH}$ e) Deconvoluted base-line corrected FTIR spectra of the amide I region showing secondary structural components of WTSOD1$^{2SH}$, ApoSOD1$^{2SH}$, ApoSOD1$^{2SH}$ (top to bottom) with excess Cu and Zn respectively. Cumulative disorder (highlighted in red) is lower in WTSOD1$^{2SH}$ than ApoSOD1$^{2SH}$. Addition of Zn but not Cu lowers disorderedness in ApoSOD1$^{2SH}$. f) Deconvoluted base-line corrected FTIR spectra of the amide I region showing secondary structural components of I113TSOD1$^{2SH}$, I113TSOD1$^{2SH}$ (top to bottom) in presence of excess Cu and Zn. g) Deconvoluted base-line corrected FTIR spectra of the amide I region showing secondary structural components of G85RSOD1$^{2SH}$, G85RSOD1$^{2SH}$ in presence of excess Cu and Zn. In all spectra, disordered/extended conformation (1637-1649 cm$^{-1}$) along with loops and turns (1662-1682 cm$^{-1}$) is represented in red.

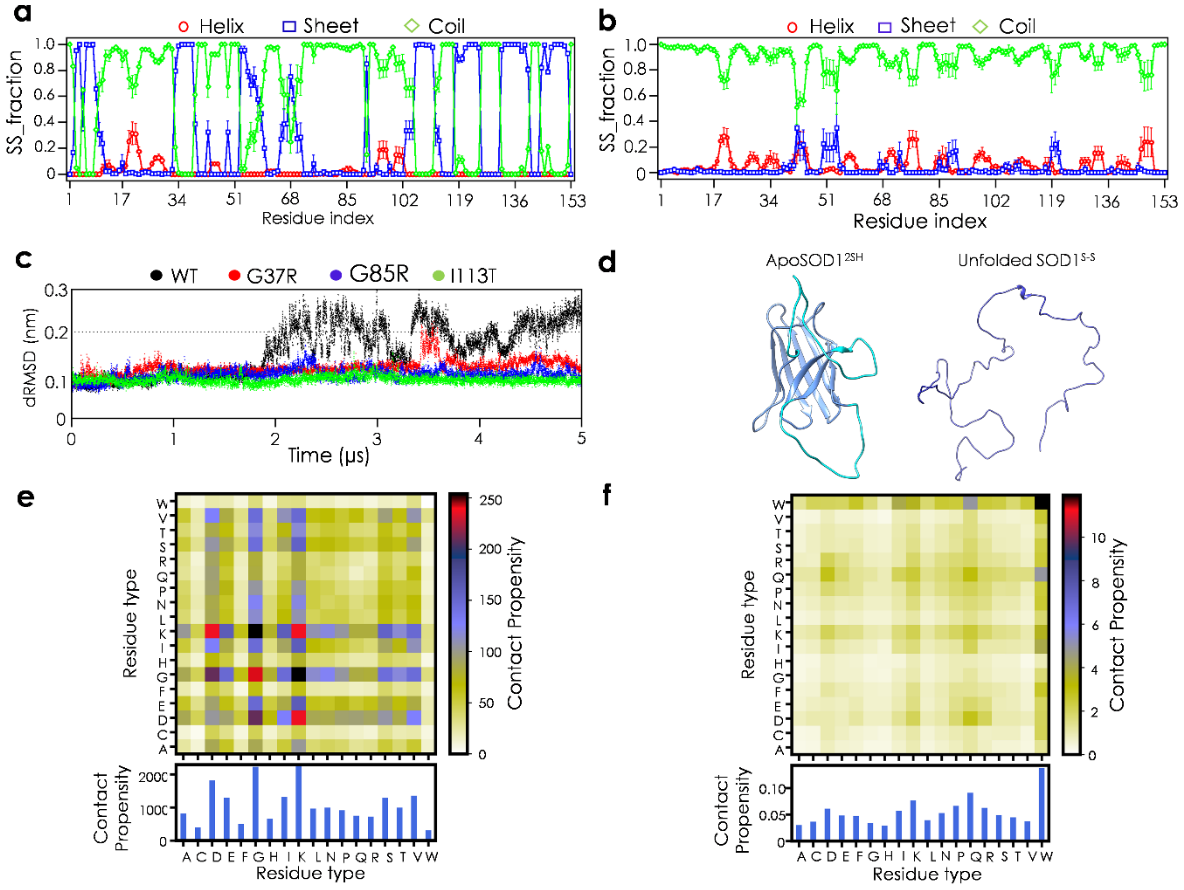


**Figure S5. Structural analysis of ApoSOD1$^{2SH}$ variants and comparison with unfolded SOD1.** a) Per-residue secondary structure analysis of ApoSOD1$^{2SH}$ b) Per-residue secondary structure analysis of unfolded SOD1. c) Cα dRMSD for β-barrel residues of ApoSOD1$^{2SH}$ variants. d)Representative snapshots of ApoSOD1$^{2SH}$ and unfolded SOD1. Time averaged intermolecular contacts within the condensed phase of SOD1 as a function of e) residue type and f) normalized by the relative abundance of each amino acid.

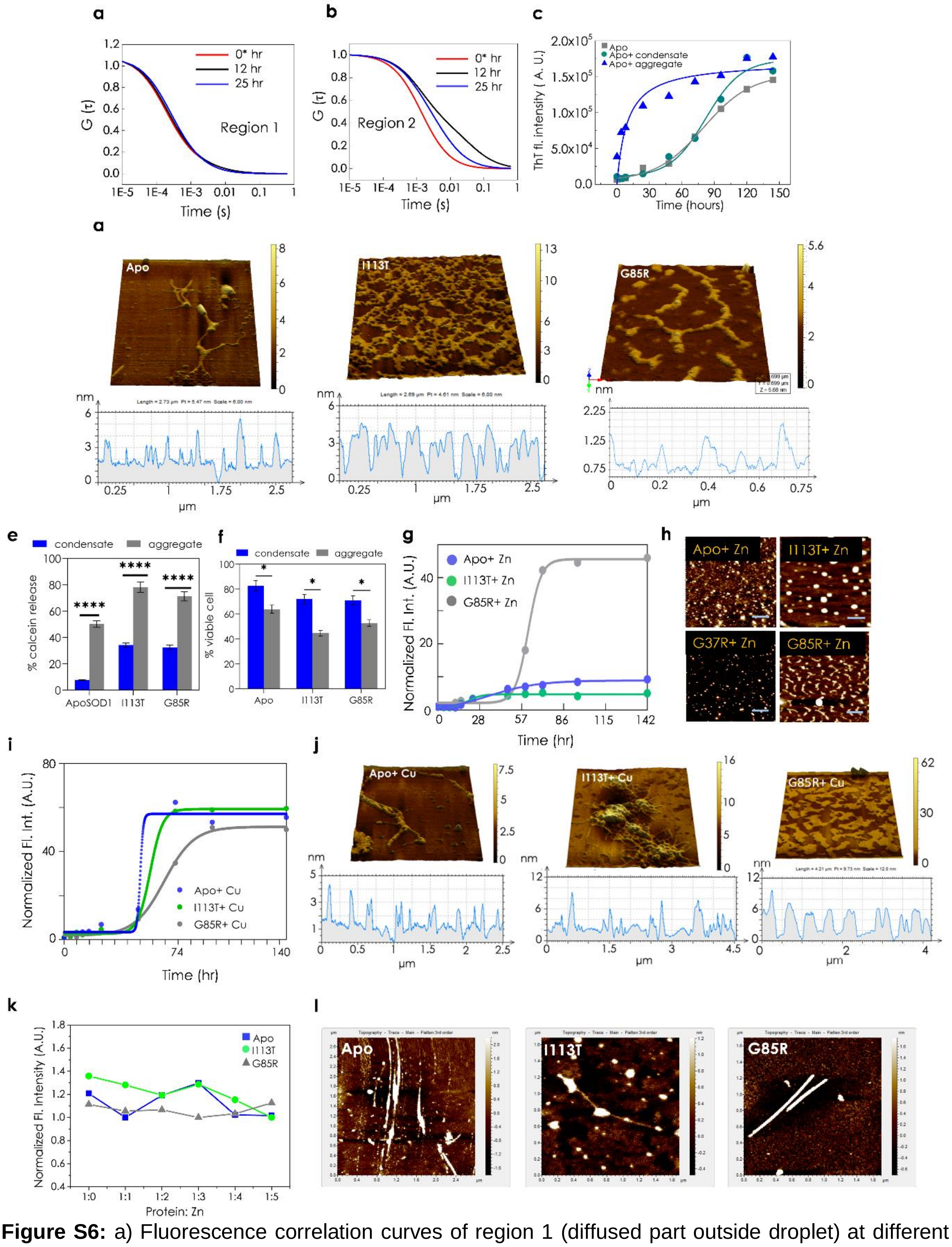

**Figure S6:** a) Fluorescence correlation curves of region 1 (diffused part outside droplet) at different time points; 0* hr is the time at which droplets form after incubation. b) Fluorescence correlation curves of region 2 (low intensity portion inside droplet) at different time points; 0* hr is the time at which droplets form after incubation. c) ThT fluorescence assay plot showing seeded aggregation of ApoSOD1$^{2SH}$ (100 µM concentration) by preformed ApoSOD1$^{2SH}$ condensates (grey) and ApoSOD1$^{2SH}$ aggregates (green) respectively as opposed to only apo protein (blue) indicate sigmoidal curves for both ApoSOD1$^{2SH}$ and ApoSOD1$^{2SH}$ seeded with ApoSOD1$^{2SH}$ condensates whereas apo seeded with 10% ApoSOD1$^{2SH}$ aggregates follow a hyperbolic kinetics. d) 3D AFM micrographs with height profiles for (from left to right) ApoSOD1$^{2SH}$, I113TSOD1$^{2SH}$, G85RSOD1$^{2SH}$ respectively. e) Calcein release assay shows that aggregates have significantly higher membrane rupture and pore forming propensities than condensates; statistical significance established using paired Student's t-test; ****$p < 0.0001$. f) MTT assay performed with SHSY5Y cell line shows SOD1$^{2SH}$ aggregates are more toxic than SOD1$^{2SH}$H condensates; statistical significance established using paired Student's t-test; *$p < 0.05$. g) ThT fluorescence assay shows aggregation kinetics of SOD1SOD1$^{2SH}$ variants incubated in presence of Zn (1:5 molar ratio) at 37 ℃, 180 RPM respectively; Zn was observed to slow down aggregation in ApoSOD1$^{2SH}$ and I113TSOD1$^{2SH}$ while no significant change in case of G85RSOD1$^{2SH}$ was seen. h) AFM micrographs of fibrillar aggregates of protein variants after prolonged incubation with Zn at 37 ℃ for 144 hours; scale bar: 200 nm. i) Aggregation kinetics of SOD1$^{2SH}$ variants incubated in presence of Cu. Cu did not have any effect on aggregation kinetics of SOD1. j) AFM micrographs of (left to right) of ApoSOD1$^{2SH}$, I113TSOD1$^{2SH}$, G85RSOD1$^{2SH}$ incubated with Cu show presence of amyloid fibrils. k) Addition of Zn to preformed SOD1$^{2SH}$ aggregates does not show fibril dissolution as observed from ThT fluorescence assay. l) AFM micrographs of preformed SOD1$^{2SH}$ aggregates (left to right) ApoSOD1$^{2SH}$, I113TSOD1$^{2SH}$, G85RSOD1$^{2SH}$ with added Zn show intact fibrils.

### 2. Supplementary tables

**Supplementary table 1.** Hydrodynamic radii of all $SOD1^{2SH}$ variants.

| | **WT** | **Apo** | **G37R** | **I113T** | **G85R** |
|---|---|---|---|---|---|
| **$r_H$ (nm)** | 2.06 ± 0.06 | 2.80 ± 0.09 | 2.13 ± 0.08 | 2.99 ± 0.07 | 2.98 ± 1.02 |

**Supplementary table 2.** Percentage of secondary structure obtained from FTIR data analysis.

| SOD1[2SH] variants | Metal ion cofactor | Cross beta (1611−1630 $cm^{-1}$) | Beta sheet (1630 -1637 $cm^{-1}$), (1682- 1689 $cm^{-1}$) | Disordered /Extended (1637-1649 $cm^{-1}$) | Alpha helix (1649−1662 $cm^{-1}$) | Disordered/ Loops and turns (1662-1682 & 1690-1695 $cm^{-1}$) |
|---|---|---|---|---|---|---|
| **WT** | **-** | 5.6±0.22 | 33.49±1.31 | - | 40.7±1.66 | 20.2±0.804 |
| **G37R** | **-** | 12±0.4 | 36.6±1.4 | - | 11.4±0.46 | 40±2.66 |
| **Apo** | **-** | 3.1±0.13 | 29.6±1.2 | 52.5±2.14 | 1±0.04 | 13.8±0.55 |
| | **Zn** | 2.7±0.12 | 33.9±1.38 | 6.9±0.28 | 49.3±2.01 | 7.2±0.29 |
| | **Cu** | 1.9±0.07 | 17.7±0.7 | 52.8±2.8 | 20.5±0.84 | 7±0.28 |
| **I113T** | **-** | 5.2±0.2 | 19.1±0.77 | 64.6±2.6 | - | 10.9±0.44 |
| | **Zn** | 14.6±0.59 | 20±0.81 | - | 57.8±2.3 | 7.6±0.3 |
| | **Cu** | 21±0.8 | 12±0.48 | 46.9±1.91 | 9±0.36 | 11±0.44 |
| **G85R** | **-** | 2±0.08 | 28±1.14 | 33±1.34 | - | 37±1.51 |
| | **Zn** | 2.6+0.1 | 31.5±1.28 | - | 1.9±0.07 | 63.9±2.6 |
| | **Cu** | 2+0.08 | 13±0.53 | - | 21±0.85 | 64±2.61 |

3. **Supplementary movie 1.** Coarse-grained (CG) phase coexistence simulation of 100 $ApoSOD1^{2SH}$ chains performed using a slab geometry. Loop IV and VII of each monomer is coloured in red and blue respectively.

### 4. Material and Methods

**Recombinant SOD1$^{2SH}$ purification**: SOD1 plasmid was transformed into competent *E. coli* (BL21 DE3 strain) cells using heat shock method. Transformed bacterial cells were grown in LB media at 37 ℃. The over-expression of SOD1$^{2SH}$ was induced with 1 M isopropyl-1-thio-β-d-galactopyranoside (SRL Diagnostics) after cells reached log phase (at $OD_{600}$ ~0.6-0.8). 1 mM $CuSO_4$ (Sigma-Aldrich) was added to the culture for proper metal loading over the protein. The cells were allowed to grow for 3.5 h after which they were pelleted down by centrifugation at 6000 rpm for 15 min at 4 °C. Cells were re-suspended in pre-chilled lysis buffer (20 mM Tris–HCl+ 500 mM NaCl, pH 8.0). After thorough re-suspension in lysis buffer, cells were sonicated (20 pulses, each of 30 s with an interval time of 1 min) to allow cell lysis. Unbroken cells and debris were removed by another centrifugation step at 12,000 rpm for 30 min. The supernatant obtained was carefully removed and allowed to bind to Ni–NTA agarose resin (Thermo Fisher Scientific, USA). The Ni–NTA column was washed using 50 ml wash buffer (20 mM Tris–HCl, 500 mM NaCl and 50 mM imidazole, pH 8.0) followed by elution with 20 mM Tris–HCl, 500 mM NaCl and 500 mM imidazole, pH 8.0. Protein concentration of eluted fractions were determined by their absorbances at 280 nm. Eluted fractions were pooled and dialysed using SnakeSkin Dialysis Tubing (10 KDa MWCO) in 20 mM Na-phosphate buffer pH 7.4. The protein concentration of the post-dialyzed fraction was estimated by recording absorbance at 280 nm using the molar extinction coefficient ($\varepsilon_{280}$= 5500 $cm^{-1}$).

**Preparation of ApoSOD1$^{2SH}$**: ApoSOD1$^{2SH}$ was prepared from the holoSOD1$^{2SH}$ by metal removal. HoloSOD1$^{2SH}$ was subjected to overnight dialysis in 50 mM Na acetate, 10 mM EDTA, pH 3.8 to carry out proper removal of metal ions. EDTA was removed by successive dialysis in 50 mM Na acetate pH 5.2 and in 20 mM Na-phosphate, pH 7.4.

**Liquid-liquid phase separation *in vitro***: To study LLPS and liquid droplet formation, 100 µM SOD1$^{2SH}$ and its mutants respectively were incubated in 20 mM HEPES buffer (pH 7.4), ~7% heparin and 100 mM NaCl for 20 minutes at 37 ℃. 10 µL of the reaction mixture was aliquoted and drop-casted onto grease free glass slides with single concavity and covered with a 22 mm coverslip (Blue Star, India), sealed with commercially available nailpolish. The slides were visualized with a 10X and 40X objective using Leica microsystems inverted fluorescence microscope in the DIC mode and fluorescence mode for imaging droplets composed of labelled protein. Droplet numbers from different fields were computed using ImageJ software. The optical density of the samples was measured at 600 nm with Hidex Sense Microplate Reader (Finland). The turbidity was measured as optical density from three independent measurements.

**Fluorescent labelling of protein**: All SOD1$^{2SH}$ protein variants were labelled using a AlexaFluor 488 Protein Labelling kit (Invitrogen) following previously established protocol (1). The fluorescence dye was dissolved in DMSO and added to 2mg/mL solution of protein under constant stirring. The molar ratio between the protein and dye was 1:10. The reaction mixture was incubated at 4℃ for 5 hours with vortexing after every 30 minutes. The labelling reaction was then quenched by adding excess β-mercaptoethanol. Excess free dye from the reaction mixture was removed by extensive dialysis in Na-phosphate (pH 7.4) buffer using SnakeSkin Dialysis Tubing (10 KDa MWCO) followed by column chromatography using a Sephadex G20 column which was pre-equilibrated with 20 mM Na-phosphate buffer (pH 7.4).

**Lifetime and Anisotropy of protein condensates:** 15 nM labelled ApoSOD1$^{2SH}$**,** I113TSOD1$^{2SH}$ and G85RSOD1$^{2SH}$ along with 100 µM unlabelled protein were subjected to LLPS conditions and droplets were visualized under a confocal microscope for Time-Domain FLIM experiments. The time-resolved fluorescence measurements were made using a time-correlated single photon counting (TCSPC) setup of Alba (ISS Inc., Champaign, Illinois). Measurement samples were excited using a 488 nm QuixX picosecond pulsed laser made by Omicorn-Laserage Laserprodukte GmbH. The repetition rate of the laser was set to be 20MHz. The laser was linearly polarized in the vertical direction, and a linear polarization cleanup filter (DPM-100-VIS by Meadowlark Optics) was used for further improving the extinction ratio. For FLIM measurements, the fluorescence emission was detected by a single photon avalanche diode (SPAD) detector (SPD-100-CTC by Micro Photon Devices) after the 530/43-nm band-pass filter (Semrock). For the anisotropy measurements, the fluorescence emission was separated by the polarization beamsplitter into two channels which are parallel and perpendicular to the orientation of the linear polarization of the excitation; both parallel and perpendicular emissions were simultaneously detected by two SPAD detectors using the same 530/43-nm band-pass filters. Both the FLIM and the anisotropy data were analyzed using the ISS 64-bit VistaVision software. For FLIM data analysis, the software allows the single or multiexponential curve fittings on a pixel-by-pixel basis using a weighted least-squares numerical approach (2-4). The single-exponential model was used for fitting the lifetime data of diffused and condensed ApoSOD1$^{2SH}$ with the instrument response function, estimated by taking the first derivative of the rising of the decay. For time-resolved anisotropy data analysis, the software performs the global fittings of both fluorescence lifetimes and rotation times. The steady-state anisotropy (*r*) was calculated by

$$r = (I_{par} - I_{per}) / (I_{par} + 2I_{per}) \quad (1)$$

where $I_{par}$ and $I_{per}$ are the measured intensities in the parallel and perpendicular channels, respectively. **Fluorescence Correlation Spectroscopy:** Fluorescence Correlation Spectroscopy (FCS) measures the fluctuations of fluorescence intensity in the confocal volume and yields the diffusion times of the fluorescent species. For nanomolar level binding study using FCS, 15nM Alexa 488 labelled monomeric protein mixed with unlabelled protein keeping total unlabelled protein (demetallated) concentration at 100 nm in presence of increasing concentration of Zn from 1:0 to 1:5 molar ratio and 10 mM TCEP. The FCS measurements were carried out using an ISS Alba FFS/FLIM confocal system (Champaign, IL, USA), coupled to a Nikon Ti2U microscope equipped with the Nikon CFI PlanApo 60X / 1.2NA water immersion objective. The 488-nm picosecond pulsed diode laser was used for the excitation for the FCS measurements. The fluorescence emission was collected using a pair of SPAD (Single Photon Avalanche Detector) detectors with the 50/50 beamsplitter and the 530/43-nm band-pass filter. The use of two detectors enabled us to determine single colour cross correlation functions to eliminate the artefacts given the detector after-pulsing. The FCS correlation curves were fit to the 3D Gaussian 1-component diffusion model, where the beam waists in the radial and axial dimensions were calibrated using a standard fluorescence dye in water of known diffusion rate.

The hydrodynamic radii of each species were computed from their respective diffusion coefficients (D). For comparison between hydrodynamic radii of the wild type protein and its mutants, Alexa Fluor 488 labelled monomeric protein (15 nm) and 100 nm unlabelled protein in presence of 10 mM TCEP in 20 mM Na-phosphate buffer (pH 7.4) was used. The FCS data obtained was normalized with respect to free dye following a previously published method (5).

In the 3D Gaussian diffusion model involving a single type of diffusing molecules (the 3D Gaussian 1-component model excluding the contributions of the triplet state), the correlation function G(τ) can be defined by the following equation

$$G(\tau) = 1 + \frac{1}{N}.\left(\frac{1}{1+\left(\frac{\tau}{\tau D}\right)}\right).\left(\frac{1}{\sqrt{1+S^2\left(\frac{\tau}{\tau D}\right)}}\right) \quad (2)$$

where $\tau_D$ denotes the diffusion time of the diffusing molecules, N is the average number of molecules within the observation volume, and S is the structural parameter that defines the ratio between the radius and the height. The value of $\tau_D$ obtained by fitting the correlation function is related to the diffusion coefficient (D) of a molecule by the following equation

$$\tau_D = \omega^2/4D \quad (3)$$

where ω is the size of the observation volume. From here, the value of the hydrodynamic radius ($r_H$) of the protein molecule/ complex/ aggregate can be obtained from D using the Stokes− Einstein formula

$$D = kT/6\pi\eta r_H \quad (4)$$

where η is the viscosity, T is the absolute temperature and k is the Boltzmann constant.

**All-atom MD simulation protocol and analysis:** Initial structure for all ApoSOD1$^{2SH}$ monomers were taken from PDB 2C9V (Chain F). Structural mutations were made through rotamer substitutions using the Dunbrack rotamer library in UCSF Chimera (6). Initial conformation for the all-atom simulation of unfolded SOD1 was taken from a 1 μs single-chain simulation performed using the single bead per-residue, HPS-Urry model in LAMMPS. The coarse-grained struture was converted to an all-atom model using MODELLER (7). All systems were modelled based on the AMBER99SB-disp force field (8) along with a modified version of the TIP4P-D water model (9). Force field parameters for Zn and Zn-coordinated residues were obtained from previous studies (10). Energy minimisation and equilibriation were performed using GROMACS 2020 (11). The SOD1 monomer was placed into an octahedral box of 6.5 nm length. Energy minimisation of the protein was first performed in vacuum using the steepest descent algorithm. Following in vacuuo minimisation, TIP4P-D water molecules were added and the solvated system was further minimised using the steepest descent algorithm. To mimic physiological salt-concentration (0.1 M), $Na^+$ and $Cl^-$ ions were added along additional $Na^+$ counter ions to achieve electrical neutrality. NVT equilibriation was performed using Noose-hoover thermostat ($T_c$ = 1.0 ps) to stabilise the system temperature at 300 K. NPT equilibriation was performed using the Berendsen barostat (12) with isotropic coupling ($T_p$ = 5.0 ps) to achieve a system pressure of 1 bar. All production simulations were performed using OpenMM 7.5 in the canonical ensemble at 300 K using the langevin middle integrator with a friction coefficient of 1 $ps^{-1}$. Masses of all hydrogen atoms were increased to 1.5 amu which allowed for a simulation timestep of 4 fs. Constraints were applied to all hydrogen-containing bonds using the SHAKE algorithm (13). Short-range non-bonded interactions were calculated based on a cutoff radius of 0.9 nm. Long-range electrostatic interactions were treated using the PME method (14). Hydrodynamic radii were calculated using the HullRad algorithm (15). All other analysis was carried out using analysis programs available within GROMACS. Per-residue root mean square fluctuation (RMSF) and distance root mean square deviation (dRMSD) were calculated using gmx rmsf and gmx rmsdist respectively. Minimum distances in G85R $ApoSOD1^{2SH}$ was measured between arginine Nη atoms and aspartate Oδ atoms using gmx mindist. Secondary structure fractions were calculated based on the DSSP library (16) using gmx do_dssp.

**CG MD slab simulation protocol:** Coexistence simulation of partially folded SOD1 were conducted using the HOOMD-Blue 2.9.3 software package (17), following the protocol proposed in our previous work (18, 19). To simulate the partially folded sequence, Loop IV and VII were constrained using the hoomd.md.constrain.rigid function (20, 21) as a separate rigid bodies, keeping the initial structure from the PDB. The initial slab configuration (15nmx15nmx280nm) was prepared from the 100 chains of SOD1 sequence based on the HPS-Urry model (22). With that slab configuration, 5μs NVT simulation (time step is set to 10 fs, Langevin thermostat friction factor, $\gamma = \frac{m_{AA}}{\tau}$, where $m_{AA}$ is the mass of each amino acid bead, $\tau$ is the damping factor, which set to 1000 ps) were conducted at 275K and 1 μs trajectory was skipped during the density profile calculation for the equilibration reason.

**Time-based maturation study using Fluorescence Correlation Spectroscopy:** 100 μM unlabelled $ApoSOD1^{2SH}$ with 15nM Alexa-488 labelled $ApoSOD1^{2SH}$ was incubated at 37℃ ,180 rpm for maturation study for 28 hours. 10μL of the sample aliquoted from different timepoints 30 mins post incubation (marked as 0*), 12 hours and ~25 hours were drop casted in a depression slide (Blue Star, India) and mounted with 22 mm coverslip (Bluestar, India) and sealed with commercially available nailpolish. The slide was visualized under confocal microscope using TD-FLIM (Nikon Eclipse Ti2, Japan) at 60X water emersion objective. FCS measurements were taken from regions with varying intensities within the droplet using point FCS mode. The FCS correlation curves were fit to the 3D Gaussian 1-component diffusion model, where the beam waists in the radial and axial dimensions were calibrated using a standard fluorescence dye in water of known diffusion rate.

**Thioflavin T (ThT) assay**: For monitoring effect of Zn on the aggregation kinetics of $SOD1^{2SH}$, all protein variants were incubated in presence and absence of Zn under shaking at 180 rpm at 37 °C for 350 hours. The protein concentrations for the aggregate preparation were 100 μM in 20 mM HEPES buffer at pH 7.4. Aliquots of proteins were withdrawn at each time-point in the aggregation pathway and diluted in 20 mM HEPES buffer at pH 7.4 to reach a volume of 500 uL. ThT was added to protein in 1:10 molar ratio (protein: ThT). Steady state fluorescence measurements were recorded in a quartz cuvette of path length 1 cm using a Photon Technology International (PTI) fluorescence spectrometer with excitation wavelength of 450 nm, emission wavelength of 485 nm and an integration time of 0.1 s averaging over 3 times. For seeded aggregation, 10% (10 μM) seed of prepared condensates and aggregates were added to 100 μM protein respectively and studied for aggregation by detecting ThT fluroscence as mentioned previously.

**Atomic Force Microscopy**: Aggregating samples of $SOD1^{2SH}$ and its mutants were aliquoted after prolonged incubation at 37°C and diluted 20 times with milliQ water. 5 μl diluted sample was drop casted on freshly cleaved mica. The aggregates were rinsed with MilliQ water and then dried using a stream of nitrogen. Images were acquired at room temperature using a Bioscope Catalyst AFM (Bruker Corporation, Billerica, MA) with silicon probes. The standard tapping mode was used to image the morphology of aggregates. The nominal spring constant of the cantilever was kept at 20–80 N/m. The spring constant was calibrated by a thermal tuning method. A standard scan rate of 0.5 Hz with 512 samples per line 6 was used for imaging the samples. A single third order flattening of height images with a low pass filter was done followed by section analysis to determine the dimensions of aggregates.

**Calcein release assay**: Calcein-loaded SUVs of the composition of 3: 7 POPC: DOPS were added to SOD1$^{2SH}$ condensates and aggregates at a protein: lipid ratio of 1:10. Lipids were purchased from Avanti Polar Lipids, USA. Within cells, the extracellular leaflet of the plasma membrane is composed of neutral phosphatidylcholine (PC) lipids such as 1-palmitoyl-2-oleoyl-sn-glycero-3-phosphocholine (POPC) or 1,2-dioleoyl-sn-glycero-3-phosphocholine (DOPC) and some sphingolipids. The intracellular leaflet is rich in negatively charged phosphatidylserine (PS) lipids such as 1-palmitoyl-2-oleoyl-sn-glycero-3-phospho-Lserine (POPS) and 1,2-dioleoyl-sn-glycero-3-phospho-L-serine (DOPS). Thus, we used POPC and DOPS for our experiments in the physiologically-relevant ratio of 3:7 PC:PS. 50 mM calcein when encapsulated within the SUVs is self-quenched, and hence shows a basal fluorescence at 515 nm when excited at 490 nm. 1 µl Triton X-100 was used to determine 100% calcein release, and all results were normalized to this value. For vesicle formation, a 3:7 ratio of POPC: DOPS was dissolved in 1 ml chloroform, followed by evaporation of the solvent under a stream of N2 gas. The resulting lipid film was hydrated in 20 mM sodium phosphate buffer, pH 7.4 containing 50 mM calcein dye. The lipid-calcein suspension was sonicated in a glass tube in the dark at 40% amplitude for 30 minutes with 30 second pulse on and 1 minute pulse off at room temperature until the sample was transparent yellow in colour. The SUVs were isolated from free dye by dialysing them in the dark in 20 mM sodium phosphate buffer, pH 7.4.

**Cell culture and cytotoxicity assay**: Neuroblastoma cell lines SHSY5Y acquired from the national cell repository (National Centre for Cell Science, Pune, India) were authenticated using STR analysis. The cells tested negative for mycoplasma contamination as tested using PCR. Cells were maintained in Dulbecco's modified Eagle's media (DMEM), which in turn were supplemented with 10% heat-inactivated fetal bovine serum (FBS), respectively, 4.5 g/L of glucose, 1.5 g/L sodium bicarbonate, 110 mg/L sodium pyruvate, 4 mM l-glutamine, 50 units/ml penicillin G, and 50 µg/ml streptomycin in humidified air containing 5% $CO_2$ at 37°C. Sub-culturing was done by allowing the passaging of cells as per ATCC recommendations (ATCC, Manassus, VA). Cells were cultured in both serum and antibiotic free culture medium before each experiment. MTT assay was employed to evaluate the cell cytotoxicity. For the initial screening experiment, the SHSY5Y cells ($4 \times 10^3$ cells per well) were seeded in a 96 well plate and incubated at 37 °C followed by treatment with different protein condensates and aggregates variants (5 µM) for 12 hr. After 12 hr of incubation, cells were washed with PBS, and then the MTT solution was added to each well and kept in incubator for 4 hrs to form formazan salt. The formazan salt was solubilized using DMSO, and absorbance was measured at 595 nm using an ELISA reader (Emax, Molecular Device).

**Fourier-transform infrared spectroscopy (FTIR)**: FTIR spectra of ApoSOD1$^{2SH}$ and SOD1$^{2SH}$ mutants in the presence of increasing concentrations of zinc and copper were acquired using Bruker 600 series FTIR spectrometer. Protein samples at a concentration of 20 µM were treated with increasing concentrations of zinc sulphate and copper sulphate, from molar ratios 1:1 to 1:5 (protein: metal ion) respectively in 20 mM Na-phosphate buffer at pH 7.4 and incubated for 15 minutes at room temperature before the measurements. All the FTIR measurements were carried out in Na-phosphate buffer. The experiments were carried out in solution, and the buffer baseline was subtracted before recording each spectrum. The spectral readouts were obtained on absorbance mode with a path length of 0.01 mm following standard methodology. The deconvolution of raw spectra in the amide I region (1700–1600 $cm^{-1}$) was done using least-squares iterative curve fitting to Gaussian/Lorentzian line shapes. Peak identifications were typically carried out by using double derivatives of the FT-IR spectra, as described before (23-25). We used OriginPro 9 software for the curve fitting, second derivative analysis and other data fitting.
