## Supplementary figures and images for "Zn-dependent structural transition of SOD1 modulates its ability to undergo liquid-liquid phase separation"

### Supplementary movie1

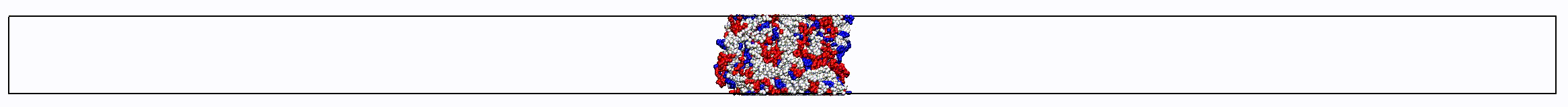
